# The neurogenic fate of the hindbrain boundaries relies on Notch-dependent asymmetric cell divisions

**DOI:** 10.1101/2021.05.31.445806

**Authors:** Covadonga F Hevia, Carolyn Engel-Pizcueta, Frederic Udina, Cristina Pujades

## Abstract

The generation of cell diversity in the central nervous system occurs during embryogenesis and requires a precise balance between cell proliferation, commitment to specific fates, and further neuronal differentiation. The cellular and molecular mechanisms regulating this balance in the embryonic brain are still poorly understood. Here we study how the neurogenic capacity in the embryonic hindbrain is spatiotemporally allocated, and how the fate and the growth of the hindbrain boundary cells are regulated. By generating a CRISPR-based knock-in zebrafish transgenic line to specifically label the hindbrain boundaries, we unveiled that boundary cells undergo a functional transition to become neurogenic during hindbrain segmentation concurrently as they maintain the progenitor cell pool. Boundary cells engaged in neurogenesis coinciding with the onset of Notch signaling, which triggered their asymmetrical cell division. Our findings reveal that distinct neurogenic phases take place during hindbrain growth and suggest that boundary cells contribute to refine the final number, identity, and proportion of neurons in the brain.

**SUMMARY:** Hindbrain boundary cells undergo a functional transition to become neurogenic concurrently as they are maintained as a progenitor cell pool. This involves a behavioral switch from symmetrically dividing progenitor cells to asymmetrically dividing progenitors, which depends on Notch-activity.

## INTRODUCTION

Tissue growth and morphogenesis are crucial interrelated processes during embryonic development, whose control is essential to generate tissues of a specific size and shape. This entails the coordinated production of different cell fates and the precise allocation of progenitor capacities over time. Tissue growth during development affects two intertwined activities: proliferation of progenitor pools leading to the increase of the number of cells, and differentiation of progenitors into post-mitotic cells (Zechner et al., 2020). In homeostatic tissues, proliferation and differentiation must be perfectly balanced to maintain a constant pool of cycling cells. However, in developing tissues, proliferation and differentiation must instead be precisely coordinated (Kicheva et al., 2014, 2012). While early on, proliferation dominates and the tissue grows, later, more cells differentiate and exit the cell cycle, and the growth rate almost ceases. In order to understand how a tissue reaches its final size it is important to know how these two processes are tuned as development progresses. Furthermore, stereotyped tissue growth must occur despite large variability in proliferation rates (He et al., 2012). Thus, balancing the rate of proliferation and differentiation in developing tissues is essential to produce organs of robust size and composition, despite the stochastic and noisy mechanisms underlying their development.

The hindbrain —the embryonic brainstem— is an interesting structure to take on this challenge, since it displays stereotypic growth dynamics with an initial phase of rapid cell proliferation, followed to a shift towards differentiation. Additionally, it undergoes morphogenesis, which establishes a crucial ground plan of regional specification. It involves the segmentation of the hindbrain leading to the transitory formation of seven metameres named rhombomeres (Kiecker and Lumsden, 2005). Rhombomeres constitute developmental units of gene expression and cell lineage compartments (Fraser et al., 1990; Jimenez-Guri et al., 2010). They are transiently visible during early development as a series of bulges in the neuroepithelium, which require the segment-restricted expression of transcription factors (for review, see Moens and Prince, 2002). They are separated by cellular interfaces called hindbrain boundaries (Lumsden and Keynes, 1989; Guthrie and Lumsden, 1991), which exhibit specific gene expression (Cheng et al., 2004; Letelier et al., 2018). Cells within these boundaries display distinct functions as development proceeds and this is beautifully coordinated with hindbrain growth (for review see Pujades, 2020). First, they act as morphomechanical barriers to prevent cell intermingling between adjacent rhombomeres to ensure that their fates and/or positional information remain segregated as they proliferate and move (Calzolari et al., 2014; Cayuso et al., 2019). Then, they become a signaling hub instructing the differentiation and organization of neurons in the neighboring rhombomeres (Cooke et al., 2005; Riley et al., 2004; Terriente et al., 2012). Concomitantly, boundary cells behave as a pool of progenitors, maintaining neural stem cells in an active proliferative state through Yap/Taz-TEAD activity (Voltes et al., 2019). Neuronal differentiation is tightly coupled to hindbrain segmentation (Schneider-Maunoury et al., 1997; Lumsden and Keynes, 1989). While rhombomeric regions are actively engaged in neurogenesis, hindbrain boundaries are devoid of proneural gene expression, acting as non-neurogenic territories (Cheng et al., 2004; Riley et al., 2004; Nikolaou et al., 2009; Voltes et al., 2019). However, recent work suggests that boundaries can provide differentiating neurons to the hindbrain (Peretz et al., 2016; Voltes et al., 2019).

Despite a good understanding of the different functions displayed by boundary cells, what their fate is, and how changes in growth, progenitor number and differentiation rates are coordinated remain elusive. Here, we have addressed these questions by reconstructing the lineage of boundary cells during hindbrain morphogenesis. Recent developments in 4D imaging (e.g., 3D plus time) and cell-tracking tools allow the assessment of cell lineages and cell behavior in the whole organ context at high spatiotemporal coverage and resolution (Dyballa et al., 2017; McDole et al., 2018; Wan et al., 2019; Wolff et al., 2018). In this work, we followed in vivo the behavior and fate of the whole boundary cell population in the hindbrain of zebrafish. For this, we have generated a versatile CRISPR-based knock-in zebrafish transgenic line that specifically targets the boundary cell population. By reconstructing the lineage of boundary cells, we unveiled that they transition from neuroepithelial stem cells to radial glia cells that undergo asymmetric divisions. Thus, boundaries constitute a self-renewing pool of stem cells and provide neurons to the hindbrain, although this neurogenic capacity is delayed in respect to the adjacent rhombomeric territories. We found that this switch in boundary cell behavior is triggered by Notch-signaling, operating through lateral inhibition along the dorsoventral axis. Eventually, boundary cells undergo differentiation, thus contributing to refine the number, identity, and proportion of neurons in the brain.

## RESULTS

### Neuron differentiation in the hindbrain boundaries is delayed with respect to the rhombomeric territories

We wanted to investigate if hindbrain boundaries were maintained as progenitor cell pools or they contributed to the neuronal lineage. Immunostaining the hindbrain with the neural progenitor cell marker Sox2 and the pan-neuronal differentiation marker HuC revealed that the neuronal differentiation domain dramatically increased over time at the expense of the progenitor domain, which concomitantly diminished (Figure 1A-1C). At 24 hours-post-fertilization (hpf), when neuronal differentiation in the hindbrain has just started, differentiated neurons were mainly allocated to the mantle zone of the rhombomeric regions (Figure 1A; see transverse view in (a)). In contrast, boundary regions did not harbor differentiated neurons at this early time point (Figure 1A; see transverse view in (a’)). Nevertheless, this changed by 32 hpf, at which point differentiated neurons were observed in boundary regions (Figure 1B; compare b-b’). By 48 hpf, there were almost no differences between the mantle zone of boundaries and neighboring rhombomeric territories (Figure 1C; compare c-c’), with both allocating many differentiated neurons. Thus, the growth of the neuronal differentiation domain in the hindbrain boundaries occurred mainly between 32 and 48 hpf, indicating a clear delay as compared to the neighboring rhombomeres. This coincided with previous studies showing that boundary cells could undergo neurogenesis when Yap/Taz-TEAD activity ended (Voltes et al., 2019). However, it could not be discerned if the differentiated neuronal pool residing in the boundaries arose exclusively from boundary progenitor cells or from the highly neurogenic neighboring regions.

**Figure 1:**
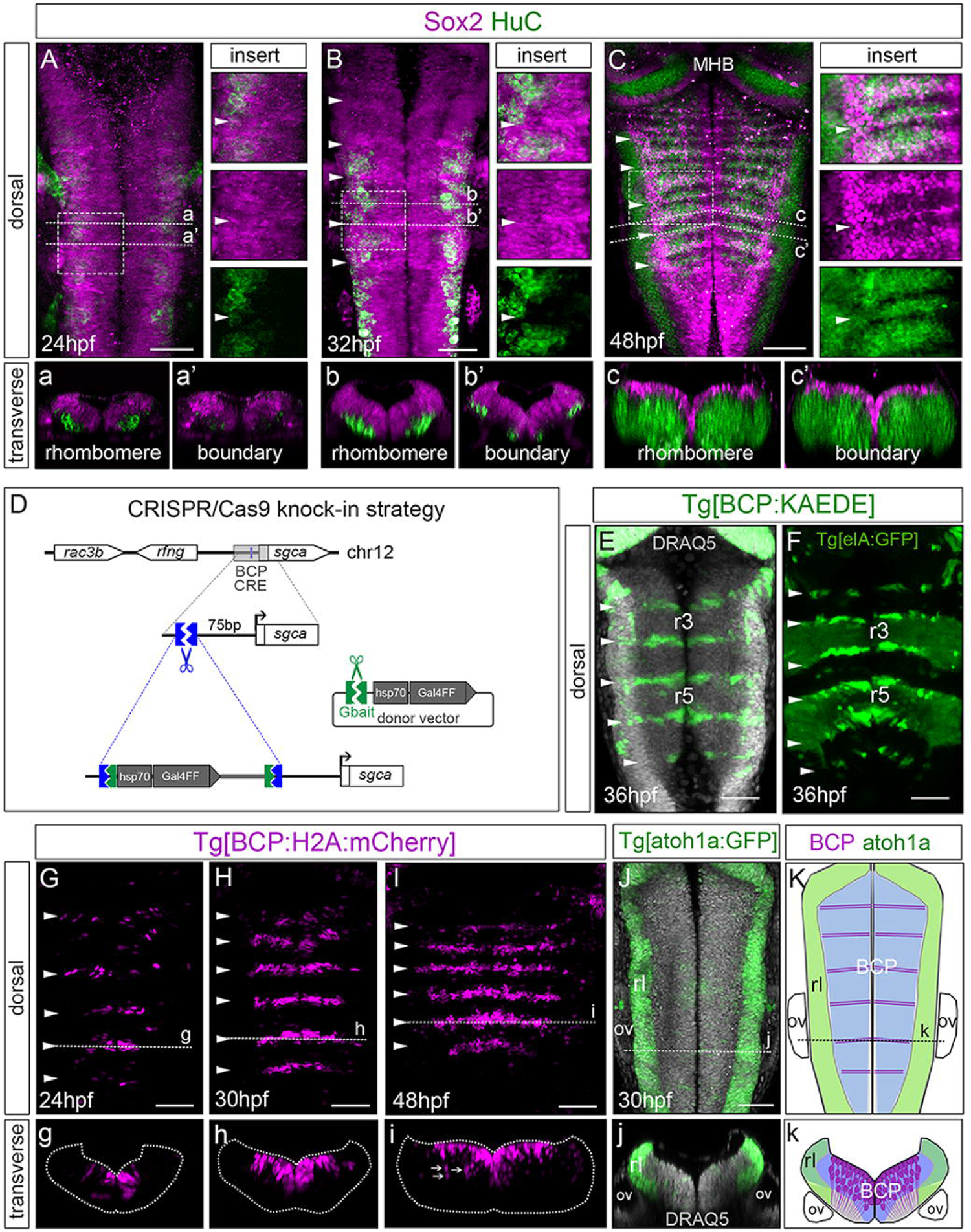
Neurogenesis in boundary cells is delayed compared to neighboring rhombomere domains. (A–C) Double immunostaining of wild-type embryos at different stages with the neural progenitor marker Sox2 (magenta), and the pan-neuronal differentiation marker HuC (green). Dorsal maximum intensity projections of the Z-stacks covering the boundary cell population are shown, with anterior to the top. Inserts, dorsal view magnifications of framed regions, with the overlay of both channels (top), only Sox2 (middle), or only HuC (bottom). Note that the boundaries display a higher accumulation of Sox2 as compared to the rhombomere-flanking regions at 32 and 48 hpf. a-c, a’-c’) transverse single-stack views at the level of rhombomeres or boundaries, respectively. Note that the HuC domain increases over time. (D) CRISPR/Cas9 knock-in strategy to generate the zebrafish Tg[BCP:Gal4] line. (E) Tg[BCP:KAEDE] embryos at 36 hpf display specific expression in the MHB and in the hindbrain boundary cells. DNA is labeled by DRAQ5 in grey. (F) Tg[BCP:KAEDE;elA:GFP] embryo at 36 hpf expressing GFP in rhombomere (r) 3 and (r) 5, and KAEDE in the boundary cells. (G–I) Spatiotemporal characterization of the Tg[BCP:H2A-mCherry] line; dorsal views are shown, with anterior to the top. g-i) transverse views of (G–I) at the level of the indicated boundary (see dashed line). White arrows in (i) indicate neuronal derivatives of boundary cells. Note that the onset of H2A-mCherry in the boundaries is around 24 hpf, and that H2A-mCherry is maintained until much later stages specifically in the boundary cells and their derivatives, excluding the rhombic lip cells. (J) Tg[atoh1a:GFP] embryo with DRAQ5, which stains the cell nuclei; (j) the transverse view is shown at the level of the indicated boundary (see dashed line). Note that GFP is restricted to the rhombic lip (Kidwell et al., 2018). (K) Scheme depicting the expression of atoh1a (green), the Sox2-domain in blue, and the boundary cells labeled by the knock-in line (magenta); k), transverse view at the level of the indicated boundary (see dashed line) showing that the domains of atoh1a and the boundary cells not fated to the rhombic lip do not overlap. Unless stated, embryos are displayed as dorsal maximum intensity projections (MIP) with anterior to the top. Transverse views as MIP of a single boundary. White arrowheads indicate the position of the hindbrain boundaries. BCP, boundary cell population; MHB, mid-hindbrain boundary; hpf, hours-post fertilization; ov, otic vesicle; rl, rhombic lip. Scale bar, 50 μ

To unveil how hindbrain boundary cells contributed to the neuronal differentiation domain during hindbrain morphogenesis, we built up a highly versatile zebrafish transgenic line that specifically labels the boundary cell population (BCP). Our strategy was based in knocking-in the Gal4 gene, into one of the cis-regulatory elements responsible for driving gene expression to the hindbrain boundary cells (Letelier et al., 2018). We targeted a genomic position located upstream of the transcriptional start site (TSS) of the sgca locus (Kimura et al., 2014) (Figures 1D, S1A–S1B); note that a target site was designed at about 90 base pairs (bp) upstream of the ATG start codon of sgca. Several transgenic lines were generated by combining the Gal4 knock-in line with different UAS reporters (Figure S1C), and efficient expression was driven to the hindbrain boundary cells (Figure 1E), as confirmed using the Tg[elA:GFP] line as a landmark for rhombomeres 3 and 5 (Figure 1F).

We generated the Tg[BCP:H2AmCherry] fish line, which expressed mCherry in the boundary cells nuclei, to study how spatiotemporally coordinated cell progenitor specification and differentiation occurred in the hindbrain boundaries alongside morphogenesis. First, we characterized this transgenic line by comparing the onset of sgca expression with the onset of mCherry. As expected, there was a short delay between the expression of the two genes: whereas sgca onset was before 20 hpf (Letelier et al., 2018; Figures S2A–S2B), mCherry was partially expressed in the boundaries at 20 hpf (Figure S2A’–S2B), while all boundary cells expressed sgca and mCherry only at 24 hpf (Figures S2C–S2D). Both sgca and mCherry expression in the boundary cells remained at least until 30 hpf (Figure S2E), as was also observed with other boundary markers such as rfng (Figures S2G–S2L). We consistently detected mCherry protein in the boundaries around 24 hpf (Figure 1G; see transverse views in (g)) and, by 30 hpf, most of the boundary progenitor cells displayed mCherry (Figure 1H; see transverse views in (h)). Interestingly, during this temporal window, we clearly observed how boundary neural progenitors, which at early embryonic stages mainly located in the medial region of the hindbrain (Figure S2a–S2i), became dorsally positioned, facing the lumen, due to hindbrain morphogenesis (Figure 1h and 1i; note that the ventricular surface occupied all the dorsal part of the hindbrain). The stability of the mCherry protein extended beyond the interval of expression of the classical boundary markers, such as rfng or sgca (compare Figure 1I with Figures S2M-S2N) (Letelier et al., 2018). This made using this transgenic line advantageous for following the fate of these progenitor cells even at later stages, in which boundary markers were no longer expressed. Accordingly, we observed mCherry-expressing cells in the neuronal differentiation domain at 48 hpf (Figures 1I and S2R), indicating that boundary cells can undergo neurogenesis (Peretz et al., 2016; Voltes et al., 2019). These results also indicated that neuronal differentiation in the boundaries did not occur at the same time than in their neighboring rhombomeric cells. As expected, the mCherry boundary cells displayed Yap/Taz-TEAD activity (Figures S2O–S2R; Voltes et al., 2019). Furthermore, the Tg[BCP:H2AmCherry] fish line was specific for boundary cells not fated to the rhombic lip, which is a transient neuroepithelial structure that expands all along the dorsolateral hindbrain and expresses the proneural gene atoh1a (Figures 1J-1K). Notably, we can use our CRISPR-based knock-in transgenic lines (Tg[BCP:reporter] lines) to specifically label the hindbrain boundary cell population and follow its behavior over time. This provides for the first time the tools to reconstruct its lineage and explore the fate of the hindbrain boundary cells.

### Boundary cells undergo neurogenesis while maintaining the cell proliferative capacity

To deeply analyze the neurogenic behavior of boundary cells, we generated double transgenic embryos Tg[BCP:H2AmCherry;HuC:GFP] and assessed the position of boundary cells and their derivatives, in both the progenitor and the differentiation domains at distinct embryonic stages. At 30 hpf, almost all boundary cells —mCherry-expressing cells— remained in the ventricular domain and did not express the differentiation marker HuC (Figure 2A, see inserts; Figure 2a; Video 1). This indicated that boundary cells were maintained as neural stem cells, and that differentiated neurons residing in the mantle zone of the boundaries at 30 hpf most probably arose from adjacent territories. However, when we performed the same analysis at 48 hpf, an average of 42% of cells expressing mCherry —e.g., derived from the boundary progenitor cell population— were in the neuronal differentiation domain and expressed HuC (Figure 2B, see inserts; Figure 2b; Video 1). This resulted in an increase of differentiated neurons deriving from the boundary progenitors during this morphogenetic period; for instance, the r5/r6 boundary had 1.6% of neurons [1.6/99 cells] at 30 hpf vs. 40.10% [79/197 cells] at 48 hpf (Figure 2C; Table 1). This demonstrates that boundary cells clearly contribute to the hindbrain by providing differentiated neurons in a non-synchronous manner with the neighboring rhombomeres.

**Figure 2:**
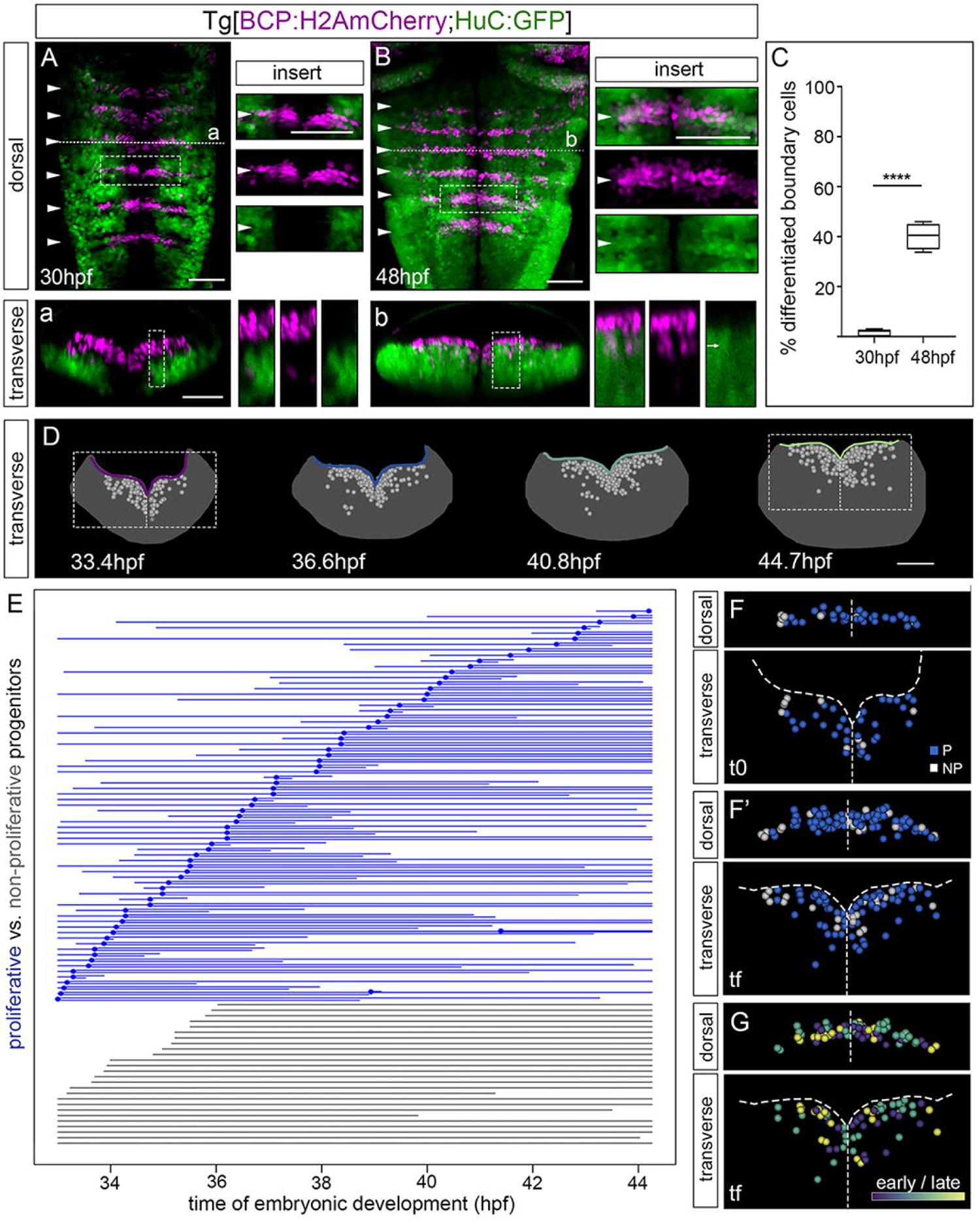
Boundary progenitor cells undergo neuronal differentiation in a time-dependent manner while they maintain their proliferative capacity. (A, B) Tg[BCP:H2A-mCherry;HuC:GFP] embryos at different developmental stages; a-b) transverse views at the level of the indicated boundary (see dashed line). Inserts (both dorsal and transverse views) display a single-stack magnification of framed regions, with the overlay of both channels (top), only boundary cells (middle), or only HuC (bottom). Note that at 30 hpf, boundary cells are in the ventricular domain and do not display HuC, whereas at 48 hpf, several boundary cells are in the neuronal differentiated domain (see white arrow in transverse view insert). (C) Boxplot showing the % of neuronal differentiated boundary cells at 30 or 48 hpf (Table 1); ****p < 0.0001. Unless stated, dorsal and transverse views are displayed as maximum intensity projections (MIP) of a single boundary. In (A, B) dorsal views anterior is at the top and white arrowheads indicate the position of the hindbrain boundaries. (D) Spatiotemporal map of a single boundary (in transverse view) with annotated nuclei from boundary cells (grey dots) displayed on the top of neural tube masks (represented as grey surfaces), showing the position of all cells at the given time. The corresponding segmented ventricular surface was color-coded according to embryonic time (33.4 hpf, purple; 36.6 hpf, blue; 40.8 hpf, turquoise; 44.7 hpf, light green). (E) Flat representation of boundary cell lineage tree in which each line corresponds to a single cell from the moment of tracking onwards that bifurcate upon cell division. An interrupted line indicates either that the movie ended or that the cell was lost from the field. The X-axis displays the time of embryonic development (hpf). Boundary cell lineages are color-coded according to proliferative behavior (blue, dividing; grey, non-dividing). (F, G) Spatial map of a single boundary (in dorsal or transverse view) displaying the regions framed in (D), with annotated nuclei from all boundary progenitors shown in (E) and displayed at t0 (33.4 hpf; F) and/or tf (44.7 hpf; F’ and G). The maps were built after following the lineages of all boundary cells during this time interval. In (F-F’), cells were color-coded according to the same criteria than in (E) and displayed at t0 (33.4 hpf) and tf (44.7 hpf). Note that different cell proliferative capacities are not specifically allocated within the boundary. (G) Spatial map of a single boundary (in dorsal or transverse view displaying the region framed in (D) with annotated nuclei from dividing progenitor cells color-coded according to time of division (33.4–36.6 hpf, purple; 36.6–40.8 hpf, turquoise; 40.8–44.7 hpf, yellow). Note that time of division does not correlate with specific spatial domains within the boundary. BCP, boundary cell population; hpf, hours-post-fertilization. Scale bar, 50 μm.

**Table 1:**
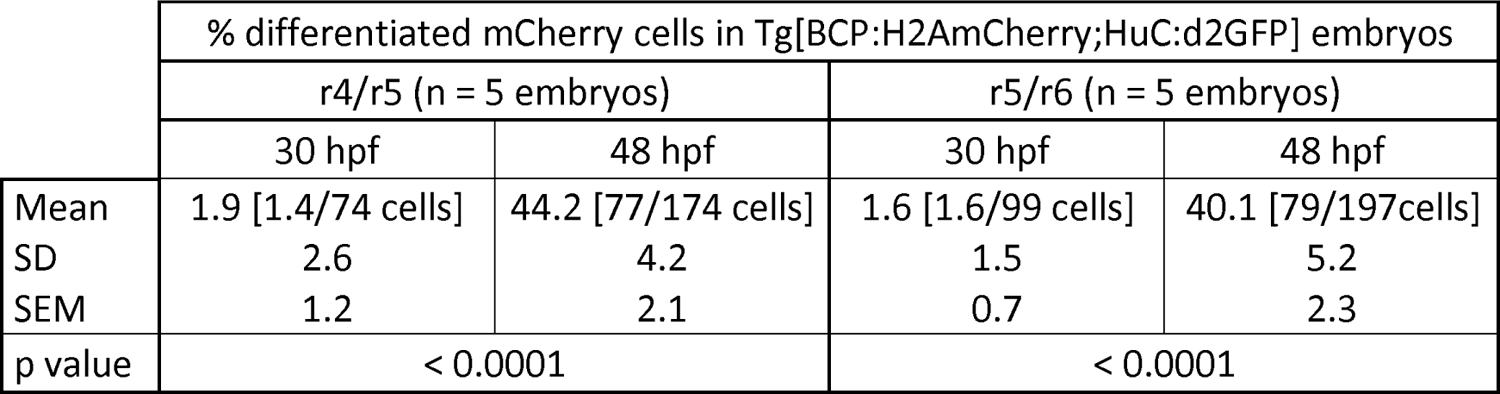
Percentage of boundary progenitor cells that underwent neuronal differentiation at different embryonic stages (****p < 0.0001).

Next question was to address how progenitor cell proliferation and differentiation are balanced in boundary cells. We explored the dynamic behavior of hindbrain boundary regions during this transition period, and gathered information on population dynamics and cell lineage relationships. For this, we made use of Tg[BCP:H2AmCherry;CAAX:GFP] and Tg[BCP:H2AmCherry;HuC:GFP] embryos, in which boundary progenitor cell nuclei and their derivatives are marked in red, and the cell plasma membranes or the differentiated neurons are marked in green, respectively. We imaged embryos with light sheet fluorescence microscopy from 32 hpf to 50 hpf (Videos 1—2), and we studied the single cell behavior of the boundary progenitors and their derivatives by tracking their position over time. This allowed us to reconstruct the lineage of boundary cells concomitantly with the assessment of the hindbrain morphogenetic changes.

First, we observed that, as the hindbrain grew and underwent morphogenesis, boundary cells continuously proliferated (Videos 1—2). The analysis of the whole boundary proliferation dynamics at single-cell resolution revealed that boundary cells did not shut down their proliferative capacity (Figure 2D—E), with a clear increase in the number of boundary cells over time (as an example, at t0 [33.4 hpf] = 103 cells vs. tf [44.7 hpf] = 155 cells). For this dynamic analysis, we made use of MaMuT software (Massive Multi-view Tracker software (Wolff et al., 2018), and cells in the boundary were depicted as dots and plotted on top of virtual hindbrain transverse sections at different time frames, with the shape of the ventricular surface color-coded according to time (Figure 2D). We analyzed the lineages of progenitor cells —those in the most ventricular domain— to assess their proliferative capacity. We considered them non-proliferative if they did not divide after following them for more than 8 h, which is the estimated average time for a hindbrain cell to divide (Lyons et al., 2003). The cell lineage tree allowed us to observe that approximately 60% of the boundary cell progenitors remaining in the ventricular domain divided during this time interval (Figure 2E; n = 73/120 cells). On the contrary, at least 26/120 cells (20%) persisted in the progenitor domain but did not proliferate during this time period, suggesting that they remained as slow- or non-proliferative progenitors. Thus, this cell lineage analysis revealed that although cells undergo differentiation, the boundary progenitor population is not depleted.

Next, we wanted to analyze whether the different cell modalities (proliferative vs. non-proliferative) were located in specific domains in the boundaries. For this, we color-coded each cell according to its proliferative capacity over the entire time period, and plotted the cells’ positions within the boundary over the entire time window (Figure 2F—F’, see maps at t0 and tf as examples). Mapping revealed that neither proliferative nor non-proliferative progenitor cells display any specific distribution in the boundary, since they were intermingled. When proliferative boundary cells color-coded according to the time they divided (early vs. late) were mapped within the boundary, we observed that progenitors were not distributed in specific domains with respect to their time of division, and that they were intermingled as well (Figure 2G). This strongly suggests that hindbrain boundaries harbor a pool of actively proliferating progenitors and at the same time they produce neurons. Moreover, it indicates that spatial and temporal features do not play a relevant role in conferring different cell behaviors in terms of proliferative capacity during this time interval.

The same analysis allowed us to observe the changes in the shape of the ventricular surface over time, revealing its flattening as the neural tube grew (Figure S3A—S3B). This change resulted from the generation of the brain ventricle (Lowery and Sive, 2009), and impacted as well the position of the progenitor cell population as previously mentioned (see better in Figure 1i and 2b). Moreover, this displacement of the boundary progenitors coincided with the increase in the neuronal differentiated population (Figure 1A—C; Video 1; Voltes et al., 2019), such that progenitors were squeezed into the space between the ventricular surface and the neuronal differentiation domain. Measuring the displacement of the whole progenitor cell population, we could observe how progenitor cells shifted their position towards the dorsal domain (Figure S3C). On the other hand, the displacement’s measurement of all tracked boundary cells along the medio-lateral (ML) axis did not result in significant changes of cell positions (Figure S3D), so we conclude that not much cell intermingling occurred along the ML axis.

### Boundary cells display different division modes

To understand how constant cell proliferation and active neurogenesis were intertwined during hindbrain boundaries growth, we explored whether the neurogenic committed cells that reside in the boundaries could emerge from the multipotent neural stem cells after division, considering the possible division modalities: symmetric, asymmetric, or a mixture of the two. For this, we followed in vivo the lineage of each single proliferative boundary cell after division and classified them according to i) the relative position of the two derivatives (besides or underneath), and ii) their location at tf, such that cells located in the ventricular zone were considered as progenitors, and cells in the ventral domain as differentiated neurons. By live tracking the boundary cells in Tg[BCP:H2AmCherry;HuC:GFP] embryos, we could observe how boundary cells underwent asymmetrical division, with one daughter cell remaining in the progenitor domain and the other undergoing neuronal differentiation, as observed by the onset of HuC expression in ventral boundary cell derivatives (Video 1). We noticed that even two adjacent proliferative progenitors could display different division modes, as depicted in Figure 3A: one gave rise to two progenitor cells (PP) by symmetric division, while the neighboring one gave rise to a progenitor and a neuron (PN) by asymmetric division. When this analysis was performed in the whole boundary cell population, we observed that proliferative progenitors displayed both modes of division during this entire time interval, with no obvious preference for symmetric or asymmetric divisions (Figure 3B; SyD, n = 24/73 cells; AsyD, n = 30/73 cells; or unclassified n = 19/73 cells, when the derivatives were not able to be tracked for a long enough time to determine their fate). Next, we investigated whether different cell division modes were allocated specifically to distinct boundary domains. Strikingly, cells displaying both division modalities intermingled within the boundary (Figure 3C— C’; see maps at t0 and tf as examples). Almost all symmetric divisions were self-renewable, giving rise to two progenitors (PP), since both daughter cells remained very dorsally located (Figure 3A, C—C’, orange cells). Thus, the cell position within the boundary was not relevant for these cell division behaviors.

**Figure 3:**
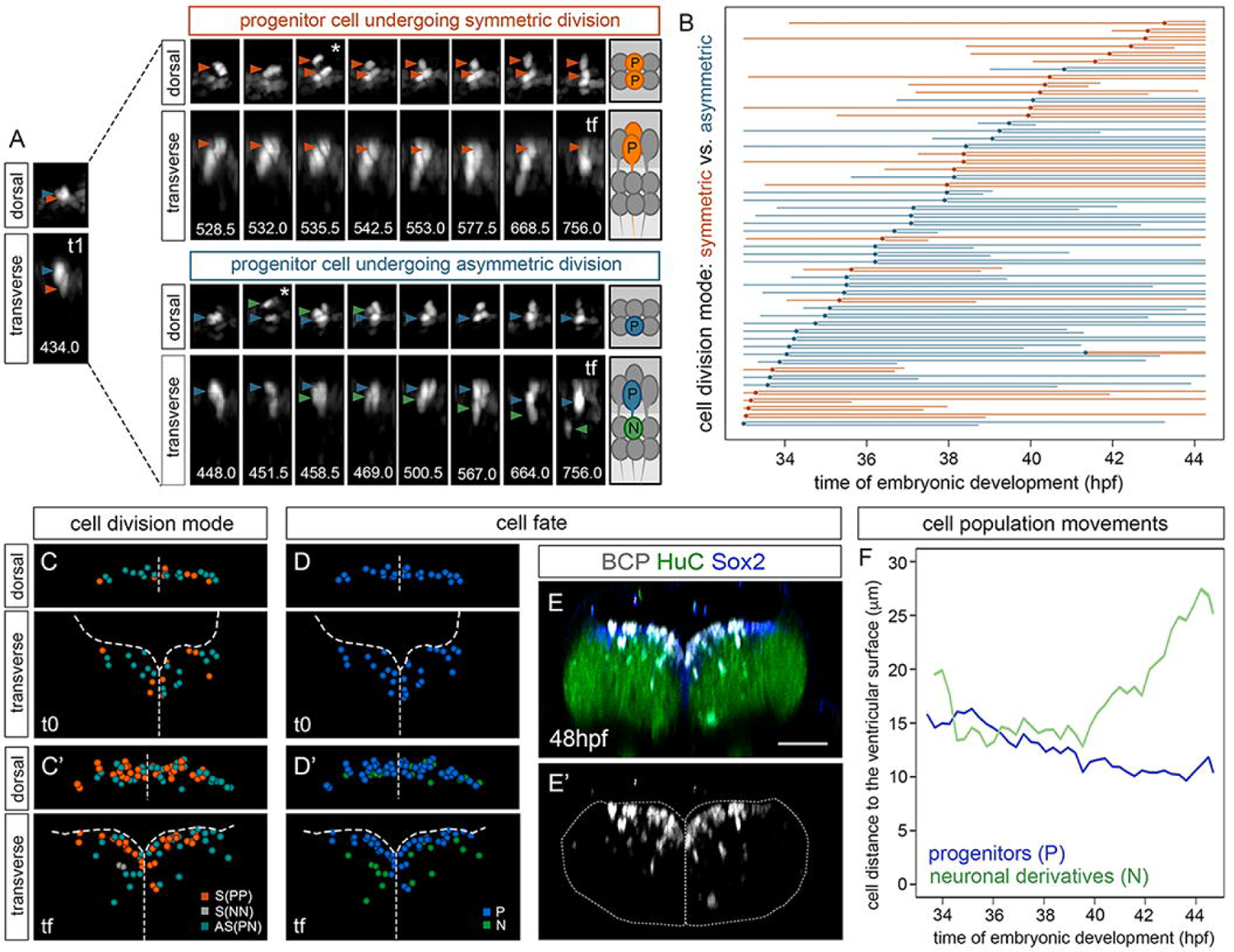
Proliferative boundary progenitor cells undergo both symmetric and asymmetric division. (A) Example of two neighboring progenitor cells (dorsal and transverse views) undergoing distinct division modes. Orange arrowheads indicate the progenitor cells that will divide symmetrically giving rise to two daughter cells remaining in the progenitor domain (P); blue arrowheads indicate the progenitor cell that will asymmetrically divide giving rise to two daughters, one progenitor (P, blue) and one neuron (N, green arrowhead); white asterisk, the time frame in which the cell divides. (B) Flat representation of proliferative boundary cell lineage tree. Each line corresponds to a single cell from the moment of tracking onwards that bifurcate upon cell division. The X-axis displays the time of embryonic development in hours-post-fertilization (hpf). Boundary cell lineages are color-coded according to mode of division (orange, symmetric; blue, asymmetric). An interrupted line means that the cell was lost from the field. Note that approximately half of the proliferative boundary cells divide symmetrically and the other half, asymmetrically. (C, D) Spatial map of a single boundary (in dorsal or transverse view displaying the regions framed in Figure 2D) with annotated nuclei from all boundary progenitors shown in (B) and color-coded according to mode of division (C) or cell fate (D). In C), orange, symmetric progenitor division; grey, symmetric neurogenic division; green: asymmetric division. Note that only two symmetric divisions were found to generate two neuronal derivatives (NN, see grey cells in the differentiated domain at tf in Figure 3C’; in both cases, the division occurred later than 40 hpf). Images were taken at t0 (33.4 hpf; C) and tf (44.7 hpf; C’). Maps were built after following the lineages of all boundary cells during this time interval. Note that different cell division modes as displayed in (C) are not specifically allocated within the boundary. See that after cell division, progenitors (blue) remain closer to the ventricular surface, whereas neuronal derivatives (green) locate in the differentiated domain (D’). (E–E’) Tg[BCP:H2AmCherry] embryos at 48 hpf and immunostained with anti-Sox2 and anti-HuC. Note that boundary progenitor cells display Sox2 and some of their derivatives express HuC. (F) Graph displaying the average distance of the two cell populations deriving from boundary cells (progenitor vs. neuron) to the ventricular surface in micrometers (0, close to the ventricular surface; 30, far from the ventricular surface) at different embryonic stages (hpf). Note how progenitor cells (blue line) become closer to the surface, and how neuronal derivatives (green line) move away from the ventricular surface. Scale bar, 50 μm.

Next, tracked boundary cells were color-coded according to their final fate and mapped them within the boundary (Figure 3D—D’). When Tg[BCP:H2AmCherry] embryos were stained for Sox2 and HuC, we detected coherent proportions of progenitors and neurons within the whole boundary cell population compared with the cell lineage analysis (compare Figure 3E—E’ with Figure 3D’). Furthermore, the asymmetric cell divisions during this period allowed the cell-renewal, and therefore the maintenance of the pool of progenitor cells within the boundaries, despite the exiting of differentiated neurons. For instance, at t0, there were 97 progenitors (43 non-dividing cells, 24 symmetrically dividing progenitors, 30 asymmetrically dividing progenitors), and at tf, there were 117 progenitors (43 non-dividing cells, 44 progenitors arisen by symmetric division, 30 progenitors arisen by asymmetric division) plus 34 differentiated neurons. To understand how these cell fate decisions impacted the overall tissue, we mapped the trajectories of the whole cell population over time by calculating the average distance of mCherry-positive cells to the ventricular surface at each time point. We observed that neurons were clearly far from the ventricular zone as they moved towards the basal domain (Figure 3F, green line), and a collective shift of the progenitor cell population towards the ventricular surface compatible with the reduction of the progenitor domain (Figure 3F, blue line). Notably, this neuronal migration occurred from 40 hpf onwards, coinciding with the time period in which we observed the gross growth of neuronal differentiated domain. This analysis provided us with information of how the boundaries function along temporal and spatial axes at the cell population level and indicated that boundary cells behaved in a non-synchronized manner: whereas some continue to divide symmetrically, others are already dividing asymmetrically and supplying neurons to the differentiated domain.

### Boundary cells undergo a Notch-dependent functional switch

Next question was to address the mechanism by which boundary cell progenitors transition towards the asymmetric division mode and neuronal differentiation, and whether the Notch-pathway played any role in this change of behavior. For this, we first assessed Notch-activity in the hindbrain using the Tg[tp1:d2GFP] transgenic line, which is a readout of Notch-active cells (Clark et al., 2012). We observed that boundary cells were devoid of Notch-activity at early embryonic stages (Figure 4A, Figure 4a, and inserts; see mCherry cells with no GFP expression). Accordingly, previous studies showed that rhombomeric cells were undergoing neurogenesis in response to Notch-signaling at early developmental stages (18—24 hpf), whereas boundary cells neither express proneural genes nor respond to Notch (Nikolaou et al., 2009). In contrast, the rhombic lip and the rhombomeric territories were Notch-active (Figures 4A; Figure 4a; Belzunce et al., 2020). However, by 48 hpf, most of the boundary progenitor cells were Notch-active (Figure 4B, Figure 4b, and inserts; see that most of the dorsal mCherry-cells facing the ventricle expressed GFP). This dramatic increase in Notch-activity was already evident before 48 hpf, as the percentage of boundary progenitor cells displaying Notch activity increased from 6.4% [2.6/40 cells] at 26 hpf to 64% [73/114 cells] to 36 hpf, as example of the r5/r6 boundary (Figure 4C; Tables 2A—2B). The detailed analysis of the Notch-active boundary cells, assessed by the Notch-reporter line, allowed us to appreciate their radial glia morphology, with the soma facing the ventricle and the radial projections towards the basal part (Figure S4A). Accordingly, these cells were positively immunostained with the zrf1 antibody (Figure S4B), a marker of radial glia processes (Trevarrow et al., 1990). Overall, the onset of Notch-activity in the boundary cells coincides with the behavioral switch of these cells to asymmetric division and towards neurogenesis, suggesting that Notch signaling is responsible for the transition of boundary neuroepithelial progenitors to radial glia cells.

**Figure 4:**
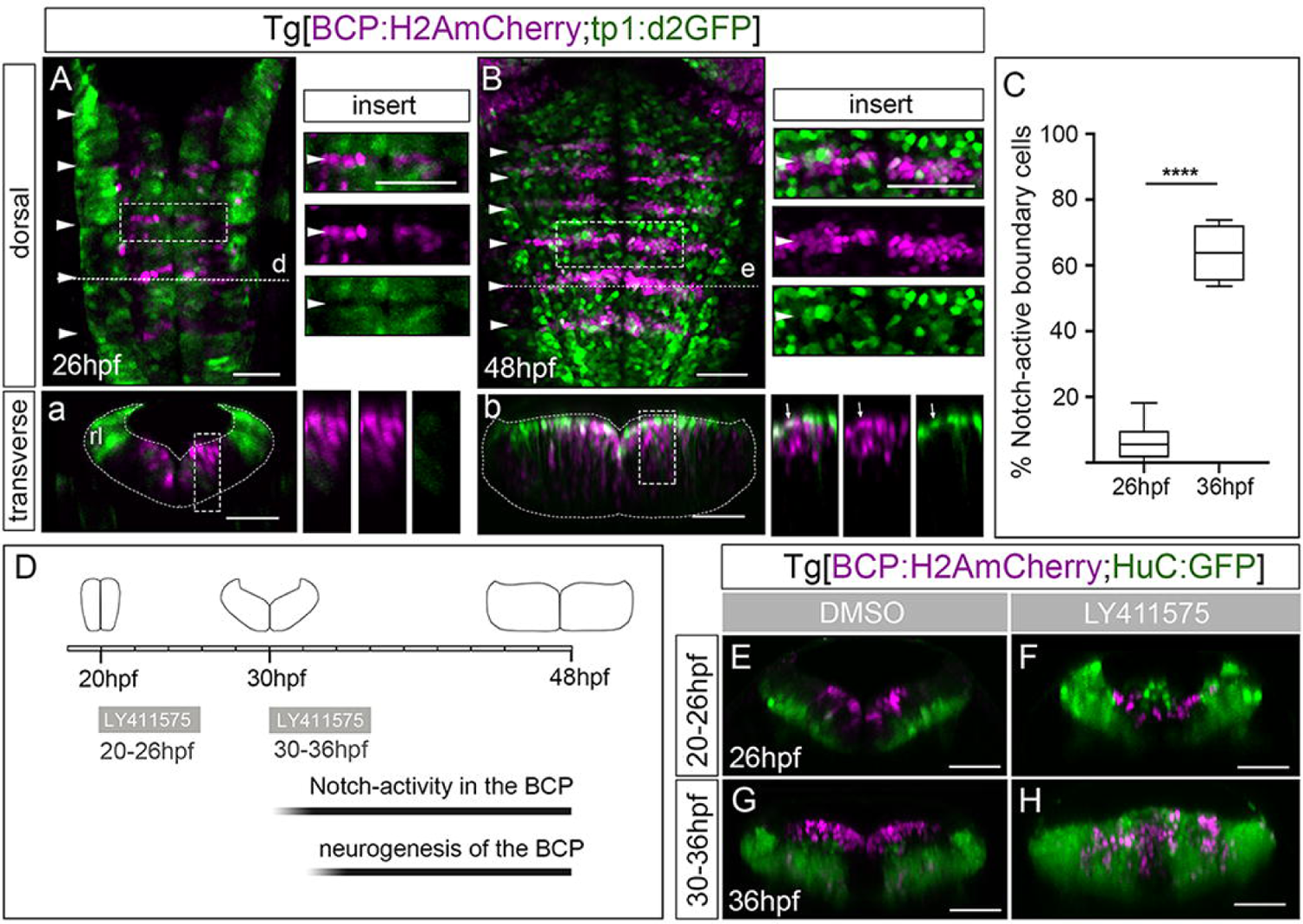
Boundary cells undergo a switch in Notch-activity. (A, B) Tg[BCP:H2A-mCherry;tp1:d2GFP] embryos at different developmental stages; a-b) transverse views at the level of the indicated boundary (see dashed line in A, B). Inserts (both dorsal and transverse views) display single-stack magnifications of framed regions, with the overlay of both channels, only boundary cells or only Notch-active cells. Note that boundary cells are devoid of Notch activity at 26 hpf but most have Notch activity at 48 hpf (see white arrow in transverse view inserts). (C) Boxplot showing the switch in Notch activity in the boundary cells (Table 2). ****p < 0.0001. (D) Experimental design for loss of Notch function experiments, depicting the shape of the hindbrain in transverse views at the distinct developmental stages. (E–H) Transverse views of Tg[BCP:H2A-mCherry;HuC:GFP] embryos treated from either 20– 26 hpf (E–F; DMSO n = 3/3 embryos; LY411575 n = 4/4 embryos) or 30–36 hpf (G–H; DMSO n = 5/5 embryos; LY411575 n = 4/4 embryos), with DMSO (E, G) or the gamma-secretase inhibitor LY411575 (F, H). For (F, H), note that boundary cells do not express HuC if treated before the onset of Notch-activity (F) but do express HuC if treated after the onset of Notch-activity (H).

**Table 2:**
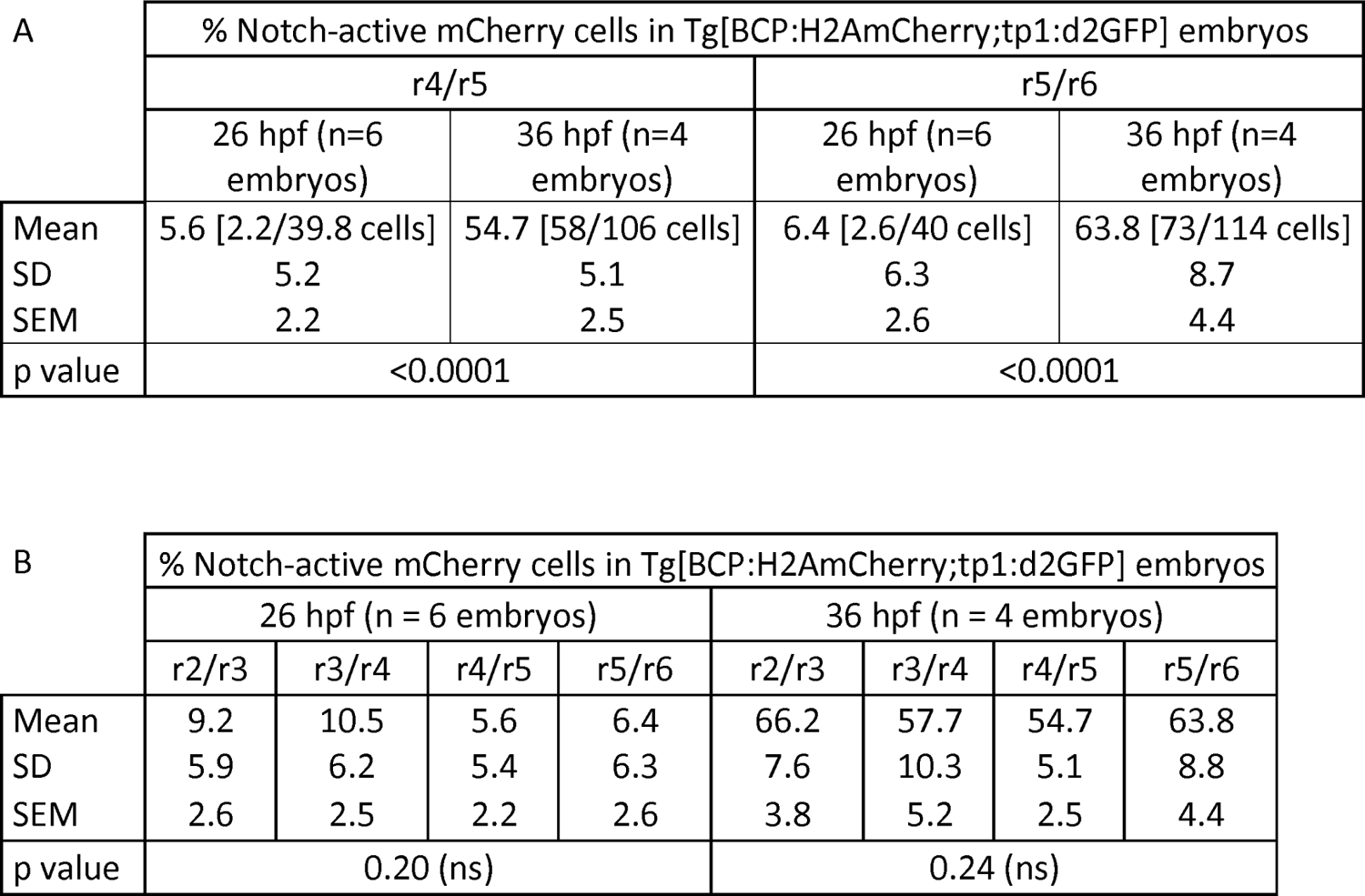
Percentage of Notch-active cells in Tg[BCP:H2AmCherry;tp1:d2GFP] embryos along the different boundaries at 26 hpf or 36 hpf (****p < 0.0001).

Then, we sought to analyze whether boundary cells responded to Notch-signaling, and assessed the effects on neuronal differentiation when Notch-activity was perturbed. We conditionally inhibited Notch-signaling in Tg[BCP:H2AmCherry;HuC:GFP] embryos by incubating them with the gamma-secretase inhibitor LY411575 at different time intervals (Figure 4D). First, we inhibited Notch when only rhombomeric regions were highly engaged in neurogenesis (20–26 hpf). As expected, the size of the HuC domain dramatically increased, although none of the mCherry-positive cells expressed HuC (Figures 4E-4F). In contrast, when Notch-activity was inhibited at the period in which boundary cells started to be engaged in neurogenesis (30–36 hpf), boundary cells massively differentiated and expressed HuC (Figures 4G—H). These results demonstrate that all boundary cells are competent for undergoing neuronal differentiation at this later stage and that Notch-activity is crucial to maintain boundary cells as radial glia progenitors.

Next, we wanted to seek the main Notch-players in the boundary cell population. When boundary cells were not undergoing neurogenesis (26 hpf), we observed a faint expression of notch3 receptor in the whole hindbrain (Figure 5A, a, and inserts; Figure S4D). Boundaries were mainly devoid of deltaD ligand at 26 hpf (Figure 5A’, a’, and inserts), although, as expected, rhombomeres expressed deltaD (Figure S4D’; Cheng et al., 2004). By 36 hpf, when approximately 64% of the boundary cells were Notch-active, all cells located at the ventricular zone displayed notch3 (Figure 5B, b, b’’; Figure S4F). Boundary cells also started to express deltaD (Figure 5B’, b’, b’’’). As expected, deltaD and notch3 expression did not overlap (Figure 5B-B’, b’’’’, and inserts). Interestingly, cells expressing notch3 were always in the most ventricular domain (Figure 5b, b’’, b’’’’), whereas deltaD-cells were just underneath notch3-cells (Figure 5b’, b’’’, b’’’’). The same dorsoventral spatial pattern was appreciated in rhombomeric regions at this stage (Figures S4D–D’; S4F–F’). This supports the previous observation that Notch-activity was restricted to the most ventricular cells, both in the boundaries and in the rhombomeres (Figure 4b; Figure S4E). The expression of notch3 receptor in the hindbrain seemed to start later than other notch receptors such as notch1a, which is already well expressed in rhombomeric cells at 18 hpf (Nikolaou et al., 2009). The fact that the increase of notch3 expression in the boundaries correlated in time with the triggering of Notch-activity in the these very same dorsal cells, and that the onset of deltaD expression occurs at the same time in the underneath cells, suggested that Notch3 and DeltaD could be relevant players in the boundary cells. Interestingly, this would imply that Notch3-DeltaD signaling would operate along the dorsoventral axis, in contrast with the salt-and-pepper distribution observed in the case of Notch1a and its ligands (Figure S4H). Then, we checked for the expression of the proneural genes ascl1b and neuroD4 as downstream targets of Notch-signaling. At 26 hpf, these genes were not expressed in boundary cell progenitors (Figure 5C—C’, c—c’’’’), although adjacent rhombomeric territories consistently expressed them (Figure 5C—C’; (Nikolaou et al., 2009). However, from 36 hpf onwards, when boundary cells engaged into neurogenesis, their derivatives expressed ascl1b and neuroD4 and, accordingly, were located in a more basal domain (Figure 5D—D’, d’’—d’’’’). These results support that the Notch canonical mechanism of lateral inhibition operates along the dorsoventral axis, regulating the balance between progenitors and differentiated cells during the neurogenic phase of the boundary cell population.

**Figure 5:**
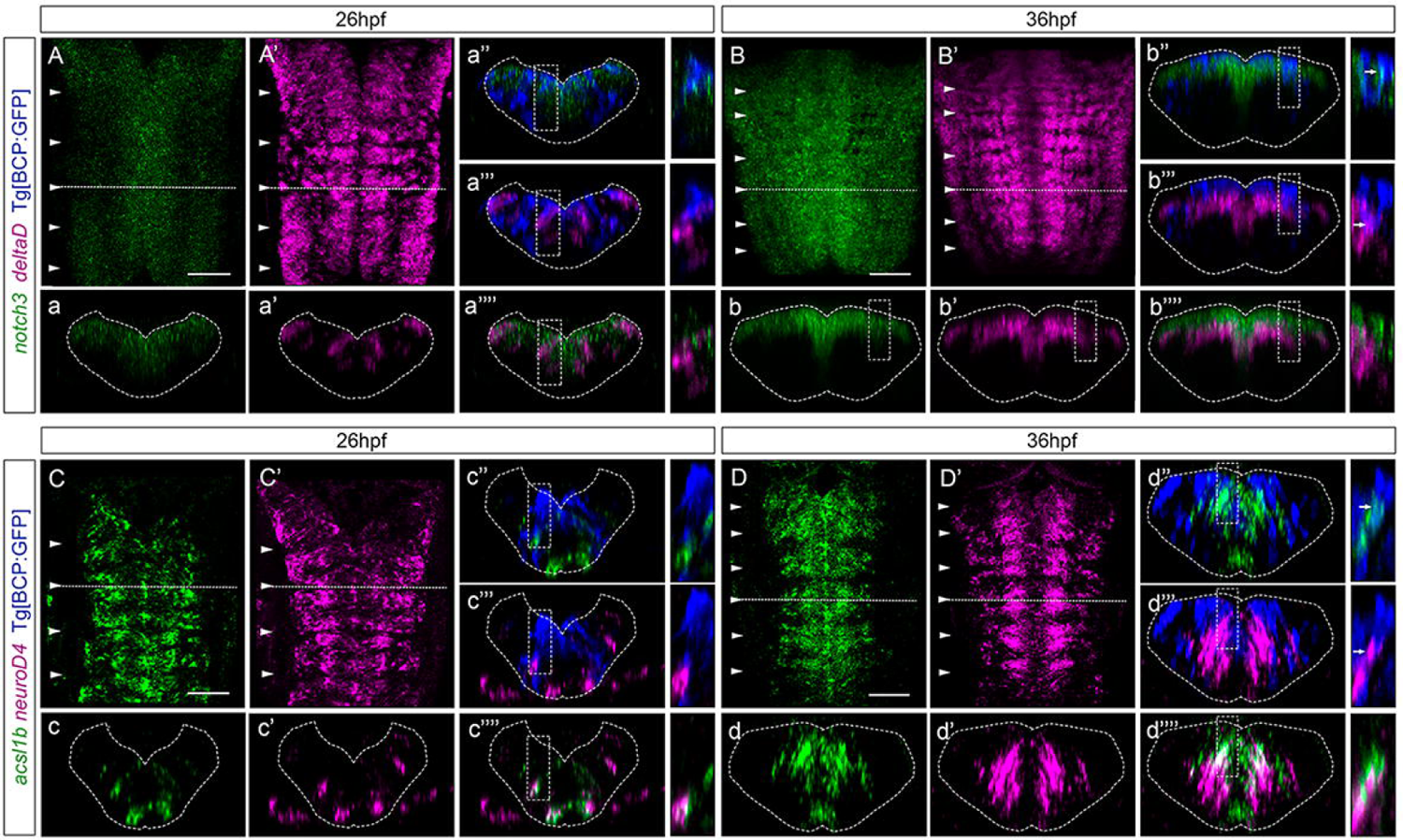
Notch players are expressed in the boundary cells from 36 hpf onwards. (A–D) Tg[BCP:GFP] embryos, before and after the onset of Notch activity in the boundary cells, were used for double in situ hybridization with: (A, B) notch3 and deltaD followed by an anti-GFP staining to detect the boundary cells. a-a’’’’, b-b’’’’), transverse views of the boundary indicated by the dashed line in (A, B), and display either single staining or several combinations according to the color-coded labeling. Inserts display magnifications of a single-stack view of the framed region. Note that notch3-positive cells are more dorsally located than deltaD-positive cells, and the stained cells never overlap; and (C-D) double in situ hybridization with ascl1a and neuroD4, followed by an anti-GFP staining to detect the boundary cells. c-c’’’’, d-d’’’’), transverse views through the indicated boundary, displaying either single staining or several combinations according to the color-coded labelling. Inserts display magnifications of a single-stack view within the framed region. Note that at 26 hpf, none of the boundary cells displayed any of the proneural genes, whereas at 36 hpf, some boundary cell–derivatives express ascl1a and neuroD4. Unless indicated, embryos are displayed as maximum intensity projections (MIP) with anterior to the top. White arrowheads indicate the position of the hindbrain boundaries. BCP, boundary cell population; hpf, hours-post-fertilization. Scale bar, 50 μm.

### Boundary cells are fated to become neurons

Boundary cells continuously proliferate combining different division modes, in such a manner that progenitor cells are maintained at the same time that they produce neurons. However, our in vivo cell lineage analyses could not inform us to the final fate of the boundary cell derivatives. Thus, to unveil whether boundary cells maintain the proliferative capacity over longer periods of time, we made use of Tg[BCP:H2AmCherry] embryos to assess the pool of boundary cells in the progenitor state much later, at 72 hpf (Figure 6A—B). The analysis of Sox2 progenitors along the hindbrain showed an enrichment in the most lateral domains with no apparent differences between rhombomeres and boundaries (Figure 6A, a), suggesting that a pool of long-lasting progenitors might reside in these lateral regions, which do not exclusively arise from boundary cells. In line with this, at this stage the neuronal differentiation domain harbored most of the boundary cells derivatives, which expressed HuC (Figure 6B, b). When we assessed the Notch-activity of the boundary progenitor cells at this stage, almost none of them displayed Notch-activity (Figure 6C, c), supporting the role of Notch-signaling in maintaining the radial glial progenitor state of boundary cells at earlier stages. In addition, we quantified the percentage of differentiated boundary cells at these two different stages of embryonic development, namely, at 48 hpf and 72 hpf (Figure 6D). We observed that an average of 33% of the boundary mCherry-cells were in the neuronal differentiation domain and expressed HuC (Table 3; [51/156 cells] as an example of r5/r6); whereas at 72 hpf, 84% of the boundary derivatives were HuC positive (Table 3, [159/189 cells] as an example of r5/r6). This percentage reflected a dramatic increase in differentiated neurons derived from the boundary progenitors at 72 hpf (Figure 6D, Table 3). This resulted in an almost depletion of progenitor cells in the boundary, as observed by the Sox2-staining (Figure 6D, see blue staining at 48 hpf but no blue staining at 72 hpf; Figure 6A), and indicated that only a small number of boundary cells remained as progenitors. This is different of what was observed in the chick hindbrain, where boundaries are highly enriched in Sox2 and can form neuorospheres in culture that self-renew and differentiate (Peretz et al., 2016). The overall number of mCherry-positive cells (boundary progenitor cells and their derivatives) did not significantly change during this time, although at 72 hpf most cells were in the neuronal differentiation domain (Table 3), suggesting that boundary cells may undergo a massive neuronal differentiation with no cell division during this last time interval (48–72 hpf). All these considered, the results indicated that the final fate of the boundary cells is to become neurons.

**Figure 6:**
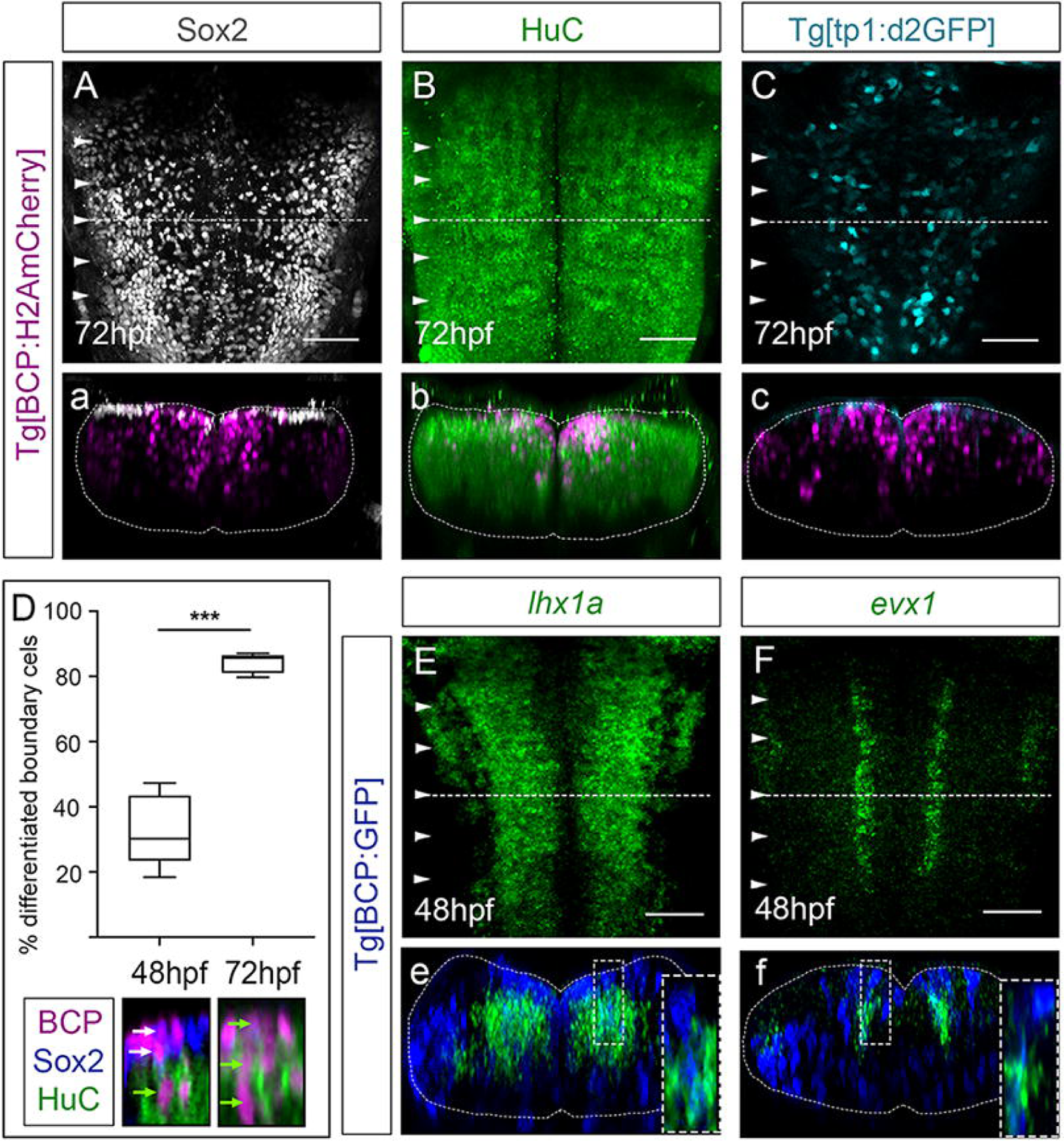
The pool of boundary progenitors extinguishes over time. (A, B) Tg[BCP:H2AmCherry] embryos at 72 hpf were immunostained with Sox2 and HuC, respectively, to assess the percentage of boundary cells that remained as neural progenitors or undergo neuronal differentiation. Note that although there are Sox2-positive cells in the hindbrain, only very few boundary cells (magenta) displayed Sox2 (white), whereas most of boundary cell derivatives express HuC (green). (C) Tg[BCP:H2AmCherry;tp1:d2GFP] embryos at 72 hpf. Note that very few boundary progenitors display Notch-activity. (D) Boxplot displaying the percentage of neuronal differentiated boundary cells at 48 hpf or 72 hpf (Table 3). Note that at 48 hpf, magenta cells located in the ventricular zone displayed Sox2 (white arrows), and those below expressed HuC (green arrows); however, at 72 hpf, very few Sox2-positive cells were observed in the ventricular zone, and most boundary cells expressed HuC (Table 3). (E, F) Tg[BCP:GFP] embryos at 48 hpf were in situ hybridized either with lhx1a (E) or evx1 (F). Inserts are magnifications of the corresponding framed regions in (E, F). In all cases, dorsal views are displayed with anterior to the top; a–c; e–f) transverse views at the level of the dotted line in (A–C, E–F). White arrowheads indicate the position of the hindbrain boundaries. BCP, boundary cell population; hpf, hours-post-fertilization. Scale bar, 50 μm.

**Table 3:** Percentage of boundary cells in Tg[BCP:H2AmCherry;HuC:GFP] embryos that underwent neuronal differentiation and the total number of boundary cells at different embryonic stages (***p < 0.001, ****p < 0.0001).

|  |  |  |  |  |
| --- | --- | --- | --- | --- |
| A | % differentiated mCherry cells in Tg[BCP:H2AmCherry] αSox2 αHuC embryos | | | |
|  | r4/r5 (n= 10 embryos) |  | r5/r6 (n=10 embryos) |  |
|  | 48 hpf | 72 hpf | 48 hpf | 72 hpf |
| Mean | 37 [54/146 cells] | 81 [136/168 cells] | 32.8 [51/156 cells] | 84.2 [159/189 cells] |
| SD | 7.8 | 6.2 | 11.1 | 3.0 |
| SEM | 3.5 | 2.8 | 4.9 | 1.3 |
| p value | <0.0001 |  | 0.0003 |  |
| B | Total number of mCherry cells in Tg[BCP:H2AmCherry] αSox2 αHuC embryos | | | |
|  | r4/r5 (n= 10 embryos) |  | r5/r6 (n= 10 embryos) |  |
|  | 48 hpf | 72 hpf | 48 hpf | 72 hpf |
| Mean | 145.6 | 168.2 | 156 | 188.8 |
| SD | 19.7 | 28.8 | 32.9 | 54.9 |
| SEM | 8.8 | 12.9 | 14.7 | 24.6 |
| p value | 0.19 (ns) |  | 0.29 (ns) |  |

To decipher whether boundary cells were fated to a particular neuronal identity, we analyzed the expression of specific neuronal differentiation markers, such as lhx1a and evx1 (Belzunce et al., 2020) in the boundary cell derivatives. We did not observe differences in the expression of these genes between neurons originated from boundaries and the rhombomeres; indeed, boundary derivatives expressed the differentiation marker that corresponded to their specific mediolateral position (Figure 6E–F). These results suggested that boundary cells are unlikely to be fated to a particular neuronal identity, but they would acquire the neuronal identity corresponding to their axial position as cells from the neighboring territories. In addition, these observations led us to propose that boundary cells eventually undergo differentiation to contribute to the refinement of the number, identity, and proportion of neurons in the hindbrain.

## DISCUSSION

Here we used cell lineage analyses to explore the behavior of hindbrain boundary cells, and the emergence of their neurogenic capacity. By high resolution live imaging over long periods of time we revealed i) the functional transition that boundary cells undergo, from progenitor cells to differentiated neurons; ii) the commitment to neurogenesis of boundary progenitors, which arises from asymmetric cell divisions that give rise to one progenitor cell and one differentiated neuron; and iii) the role of Notch-signaling in the regulation of this asymmetric division to ensure the balance between proliferating cells and cells engaging in neuronal differentiation. Disruption of the Notch-pathway –and therefore, asymmetric cell division mode– results in massive neuronal differentiation. These results now provide the cellular and molecular data – summarized in the model of Figure 7– to complement the well-described gene regulatory networks involved in neuronal differentiation.

**Figure 7:**
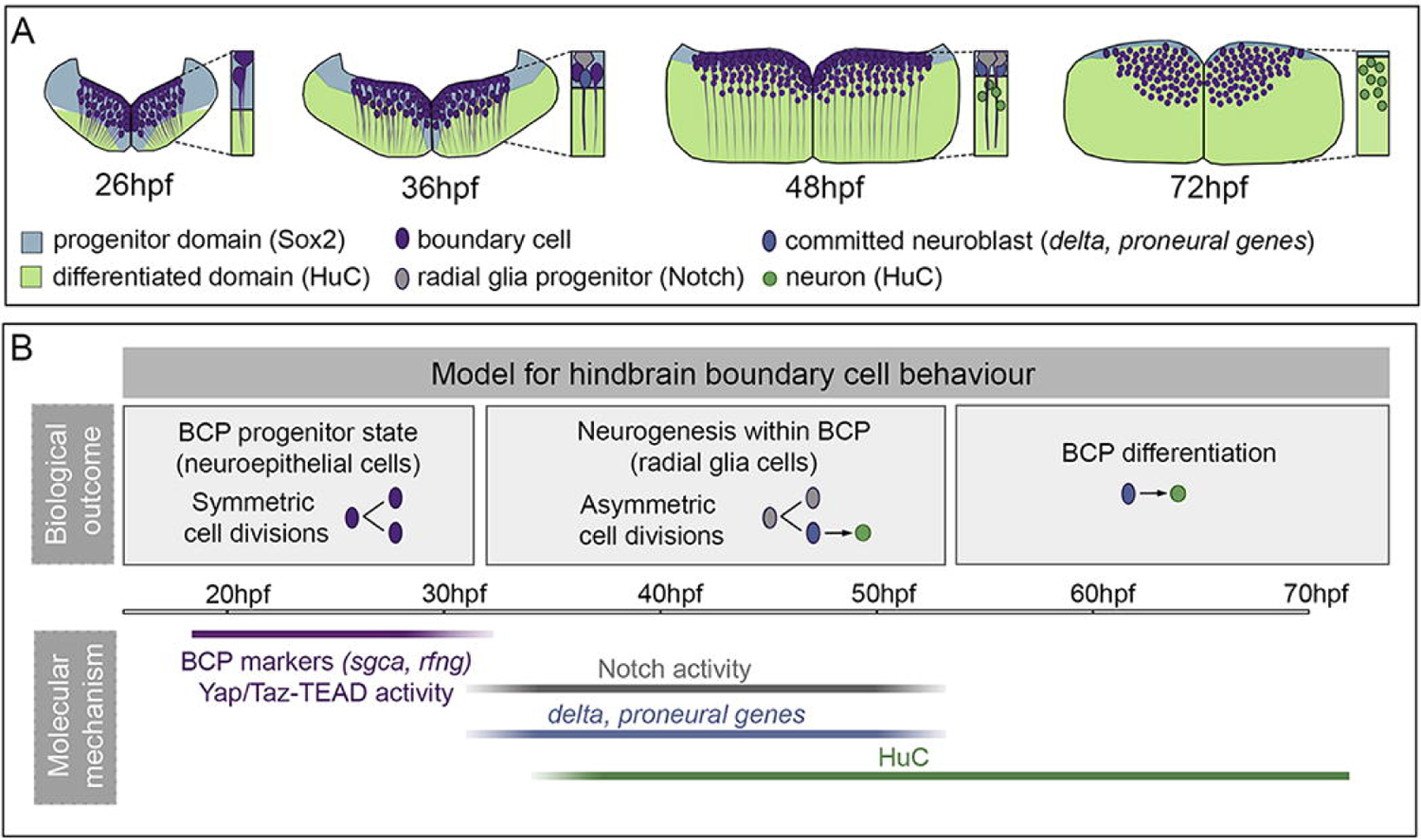
Boundary cells undergo behavioral changes during hindbrain morphogenesis. (A) Illustration depicting the functional transition of the boundary cell population. At the beginning of hindbrain neuronal differentiation (26 hpf), the cells within the boundary are neuroepithelial progenitors that divide symmetrically (purple cells). Boundary progenitor cells activate Notch-signaling around 32 hpf, singling cells out (grey cell); these singled-cells undergo asymmetric cell division (similar to radial glia cells) giving rise to one progenitor (grey cell) and one committed neuroblast (blue cell), such that the pool of boundary cells is maintained while still contributing differentiated neurons to the tissue. By 48 hpf, the progenitor domain is reduced and squeezed in between the ventricular surface and the neuronal differentiation domain; the most dorsally-located boundary progenitor cells still display Notch activity, whereas the ones committed to the neuroblast lineage differentiate into neurons (green cells). Finally, the pool of boundary progenitor cells extinguishes over time. (B) Model explaining the boundary cell behavior during this time interval, and the corresponding associated molecular features. Boundary cells are specified at the interfaces between rhombomeres at early embryonic stages (18–20 hpf) and express different boundary markers (Amoyel et al., 2005; Letelier et al., 2018; Cayuso et al., 2019). They activate Yap/Taz-TEAD in response to mechanical stimuli to be maintained as proliferating progenitors and expand the progenitor pool (Voltes et al., 2019). By 32 hpf, boundary cells become Notch-active, and they can engage in neurogenesis. This Notch switch coincides with the acquisition of a new mode of cell division — asymmetric division—, which allows boundary cells to provide differentiated neurons to the hindbrain. Accordingly, boundary derivatives display proneural genes and neuronal differentiation markers such as HuC. Finally, boundary cells most probably undergo direct differentiation, as the differentiation domain dramatically increases at the expense of the progenitor domain.

The hindbrain is a crucial coordination center in the Central Nervous System, where the neurogenic capacity is differentially allocated according to the position of the progenitor populations along the anteroposterior (AP) axis during embryogenesis. Rhombomeric cells engage very early in neurogenesis in a Notch-dependent manner (Nikolaou et al., 2009; Gonzalez-Quevedo et al., 2010), such that the first neurogenic phase in the hindbrain is contributed by rhombomeric progenitors. On the other hand, boundary cells are delayed in their neurogenic commitment, and are kept as neuroepithelial progenitors that can actively proliferate to expand the stem cell pool in a Yap/Taz-TEAD-dependent manner (Voltes et al., 2019). Here, we show that hindbrain progenitors are heterogenous with respect to Notch-signaling and their neurogenic commitment. During the early neurogenic phase, Notch is inactive in the boundary cells –when they are neuroepithelial progenitor cells dividing symmetrically–, whereas Notch is highly active in the adjacent regions remarkably engaged in neurogenesis. Later, Notch is important for cell fate acquisition in boundary cells, resulting into a switch of the boundary cell division mode. Thus, upon asymmetric division of boundary cells, the progenitor daughter cell would remain in the ventricular region as a radial glia-like progenitor displaying Notch-activity, Sox2 and zrf1. On the other hand, the daughter cell engaged in neurogenesis would migrate in axial direction towards the ventral domain, first expressing delta and proneural genes and eventually differentiating into neuron. Similar scenarios, in which neurogenic commitment arises through Notch-signaling conferring the capacity to asymmetrically divide have been described in other neural systems, such as the retina, the mammalian neocortex, and the spinal cord (Nerli et al., 2020; Mizutani et al., 2007; Kressmann et al., 2015). Thus, the most plausible explanation is that this functional switch in Notch-activity results in the transition of boundary cells from a restricted symmetric division mode to an acquired asymmetric division capacity, which is crucial to maintain the balance between proliferation and differentiation. This allows to ensure covering the neurogenic needs and to maintain the proliferative pool. Noteworthy, the mechanism of signaling in the boundaries driven by Notch3, seems to be based on lateral inhibition at the dorsoventral level, according to the spatial distribution of the Notch3 receptor and ligands. However, what triggers Notch-activity in the boundaries at that given moment remains to be unveiled. It has been suggested that the sustained rfng expression in boundaries during 16 hpf–30 hpf, promoted Notch-activation, which inhibited neuronal differentiation (Cheng et al., 2004). Our work suggests that although boundary cells may express members of the Notch-pathway, they do not display Notch-activity and they are not responsive to Notch before 30 hpf. In line with this, embryos in which Notch-pathway was disrupted early –either mid^ta52b^ mutants or embryos pharmacologically treated at 16 hpf– did not display proneural gene expression in the boundaries, demonstrating that at early stages they are not responding to Notch (Nikolaou et al., 2009).

Previous studies using single-cell clonal analyses in the whole hindbrain suggested that the majority of hindbrain neurons are born from symmetric neurogenic divisions — independently of the cell position along the AP axis— and that only a small percentage of asymmetric divisions occurs (Lyons et al., 2003). The strength of our strategy is that we can live-monitor the behavior of the entire boundary cell population; in other words, we can elucidate the emergent properties of the entire cell population. As boundary cells constitute a small proportion of hindbrain progenitors, it is possible that the clonal analyses of Lyons et al. (2003) contained a small number of boundary cells, and that the observed asymmetric divisions corresponded to boundary cells. This would suggest that whereas the main part of the hindbrain favors mainly symmetric neurogenic divisions, boundaries would behave different and be able to contribute to the growth by asymmetrically dividing. Our interpretation is that the different temporal distribution of progenitor/neurogenic capacities along the AP axis would make that boundary cells engage in neurogenesis but in a delayed manner: rhombomeric progenitor cells would be the main contributors to neurons in the first phase of neurogenesis, and boundary cells would participate later. Thus, although the different progenitor pools along the AP axis are not synchronized, they might share steps of a similar program, as is observed in other contexts for regulating the growth of the tissue: expanding the stem cell niche (symmetric self-renewing divisions), maintaining the progenitor pool (asymmetric divisions), and finally differentiating neurons (either by symmetric neurogenic divisions or terminal differentiation) (see reviews, Shimojo et al., 2008; Sueda and Kageyama, 2019). Notably, we observed that boundary cells are unlikely to provide specific neuronal identities, but that boundary derivatives acquired their neuronal identity according to the Cartesian grid they interpreted —in other words, according to their positional information. This observation, together with the fact that boundary progenitors are mostly exhausted by 72 hpf, suggests that boundary cells after asymmetric division may end up differentiating and are required for supplying neurons to adjust the final number and proportions, more than for defining new neuronal subtypes. Interestingly, it indicates that overall neurogenesis would require a less tight regulation because the system could be finally adjusted by cells deriving from adjacent regions. Therefore, boundary cells would work fine-tuning the neuronal population. Whether this proliferative capacity remains beyond 72hpf by a remaining small fraction of putative progenitors is definitively a possibility (we did not address this here due to technical constraints). However, we cannot rule out an alternative scenario in which this later phase of neurogenesis would provide different neuronal subtypes contributing to the lineage diversity, such as it happens in the cortex (for review see Beattie and Hippenmeyer, 2017). As we do not know yet how and when different neural types are produced in the hindbrain, this is still a plausible consideration.

A question that now remains to be addressed is how this diversity arises from groups of “equivalent” boundary progenitor cells. While a body of previous work had suggested an orderly deterministic program for progenitor cell fate decisions in the mammalian cortex (Gao et al., 2014), a more stochastic model is currently favored (Llorca et al., 2019). The stochastic model has also been shown in the adult zebrafish telencephalon (Than-Trong et al., 2020), and the retina (He et al., 2012; Zechner et al., 2020). Our data showed that neither space nor time foretell the further behavior of boundary cells, suggesting that cell states and cell fates may be evenly distributed in the boundaries and thereby favoring a stochastic model.

## MATERIALS AND METHODS

### Zebrafish strains

Zebrafish (Dario rerio) were treated according to the Spanish/European regulations for the handling of animals in research. All protocols were approved by the Institutional Animal Care and Use Ethic Committee (Comitè Etica en Experimentació Animal, PRBB) and the Generalitat of Catalonia (Departament de Territori i Sostenibilitat) and were implemented according to European regulations. Experiments were carried out in accordance with the principles of the 3Rs.

Embryos were obtained by mating of adult fish using standard methods. All zebrafish strains were maintained individually as inbred lines. The Tg[βactin:HRAS-EGFP] line (termed Tg[CAAX:GFP] herein) expresses GFP in the plasma membrane and was used to label the cell contours (Dale and Topczewski, 2011). The Tg[tp1:d2GFP] line is a readout of cells with Notch activity, which expresses destabilized GFP under the control of tp1 promoter (Clark et al., 2012). The Tg[HuC:GFP] line labels differentiated neurons (Park et al., 2000). Tg[atoh1a:GFP] (Belzunce et al., 2020) labeling the rhombic lip was generated by crossing Tg[atoh1a:KalTA4] (Distel et al., 2009) and Tg[UAS:GFP].

### Generation of CRISPR-Cas9 knock-in transgenic line

The Gal4FF gene was inserted in one of the cis-regulatory elements (CRE) driving gene expression to boundary cells (Letelier et al., 2018). First, three sgRNAs (#1, #2 and #3) were designed to target the cis-regulatory element (CRE) located upstream of the sgca transcription start site (Figure S1A–1B). For this, the best hits provided by crisprscan.org and chopchop.cbu.uib.no platforms were used. Three sgRNAs (#1, #2 and #3) were synthetized as described in Gagnon et al., 2014 and their efficiency was confirmed by detecting indels in the genome target site. To do so, one-cell stage embryos were injected with each sgRNA in combination with Cas9 protein, and gDNA was extracted and amplified by PCR using Fw1 (5’-GAG AGA CCA CAC AGG GAA GC-3’) and Hex-labeled Rv1 (5’-AGA GCA TAT ACA GAT CTA GGA AGA ATC-3’) primers. The PCR product was subsequently sequenced by Gene Scan (Figure S1C). The presence of a multi-peak pattern indicates indels in the genome target site. sgRNA #2 displayed the best efficiency for digesting genomic DNA (40%). Gbait-hsp70:Gal4FF described in Kimura et al., 2014 was used as donor vector and contains the eGFPbait (Gbait) as sgRNA target site, hsp70 promoter and Gal4FF (Figures 1D, S1A–S1B). For generating the Gal4FF knock-in transgenic line, Tg[UAS:KAEDE] embryos at the one-cell stage were injected with 2 nl of a mix containing 50 ng/μ of each sgRNA (sgRNA #2 for digesting genomic DNA, and Gbait sgRNA for digesting the donor plasmid), 300 ng/μ of Cas9 protein with NLS (CP01-50 PNAbio) and 12 ng/μ of phenol:chloroform purified donor vector, so that concurrent double-strand breaks in the genomic target locus and the donor DNA could result in its integration at the target locus (see Figure S1 for a more detailed description of the knock-in pipeline, screening and genotyping strategy). The Gal4FF primers used were the following: Fw 5’-GCA GGC TGA AGA AGC TGA AG-3’ and Rv 5’-GGA AGA TCA GCA GGA ACA GC-3’ resulting in 59% efficiency (n = 16/27). Knock-in founders were selected by KAEDE-expression in F1 offspring and displayed a germline transmission rate of 25% (n = 4/16). 75% of these founders specifically expressed KAEDE in the boundaries (n = 3/4). Finally, adults were genotyped by fin clip using a strategy that confirmed integration and the insert directionality (Figure S1C); this involved a PCR with the following primers: Fw1 5’-GAG AGA CCA CAC AGG GAA GC-3’; Rv1 5’-AGA GCA TAT ACA GAT CTA GGA AGA ATC-3’; Fw2 5’-ATC GCA GAC CGA TAC CAG GA-3’; and Rv2 5’-TCG GGG AAA AAG TCC TGT CA-3’. The selected Tg[BCP:Gal4] fish used in this work contained 2 copies of the donor vector inserted in tandem (forward and reverse respectively). The Tg[BCP:Gal4;UAS:KAEDE] line (called Tg[BCP:KAEDE] along the manuscript) was outcrossed to segregate the UAS:transgene and isolate the Tg[BCP:Gal4]. After this, several transgenic lines were generated by crossing Tg[BCP:Gal4] fish with different UAS:reporter (UAS:GFP, UAS:H2AmCherry; Figure S1C). Specific expression was efficiently driven to the hindbrain boundaries (Figure 1E-1F). Certain leaky activity was appreciated in some ventrolateral cells in both boundaries and rhombomeres when Tg[BCP:Gal4] line was combined with UAS:KAEDE and UAS:GFP reporters, but not with UAS:H2AmCherry.

### Whole mount *in situ* hybridization

Embryo wholemount in situ hybridization was adapted from Thisse and Thisse (2008). The following riboprobes were generated by in vitro transcription from cloned cDNAs: deltaD (Haddon et al., 1998) and neurod4 (Park et al., 2003). KAEDE, mCherry, and notch3 probes were generated by PCR amplification adding the T7 promoter sequence to the Rv primer: KAEDE Fw 5’-GCT GCT TAT GGA AGG CAA TG-3’ and Rv 5’-CAA TCC AGA ATG GGC AAC AG-3’; mCherry Fw 5’-CCA AGC TGA AGG TGA CCA AG-3’ and Rv 5’-GCT CGT ACT GTT CCA CGA TG-3’; notch3 Fw 5’-ATG GGG AAT TAC AGC CTT TG-3’ and Rv 5’-GGC AAA CAA GCA CAT TCG TA-3’. sgca and rfng probes were generated as described (Letelier et al., 2018), ascl1, lhx1a and evx1 as described (Belzunce et al., 2020), and notch1a as described (Taberner et al., 2020). For fluorescent in situ hybridization, FLUO- and DIG-labeled probes were detected with TSA Fluorescein and Cy3, respectively.

### *In toto* embryo immunostainings

For immunostaining, 4% PFA-fixed embryos were permeabilized with 10 mg/ml proteinase K for different times according to the developmental stage (24–28 hpf, 5 min; 30–36 hpf, 10 min; 48–56 hpf, 20 min; 72 hpf, 30 min), blocked in 2% BSA, 5% goat serum, 0.1% Tween20 in PBS for 3 h at room temperature, and then incubated overnight at 4°C with the primary antibody. The primary antibodies used were: rabbit anti-GFP (1:500; Origene TP401), mouse anti-GFP (1:500; ThermoFisher A-11120), rabbit anti-Sox2 (1:100; Abcam ab97959), mouse anti-HuC (1:400, ThermoFisher A-21271), rabbit anti-Taz (1:150; Cell Signaling, D24E4), mouse anti-ZO1 (1:500 ThermoFisher ZO1-1A12), and zrf1 (ZDB-ATB-081002-46, ZIRC). After extensive washing with PBST, embryos were incubated overnight at 4°C with secondary antibodies conjugated with Alexa Fluor 488, 594, or 633 (1:500; Invitrogen). DRAQ5 (1:2000; Biostatus, DR50200) was used to stain nuclei.

### Confocal imaging of whole mount embryos

Anesthetized live embryos expressing genetically encoded fluorescence or stained fixed samples were mounted in 1% low melting point (LMP) agarose with the hindbrain positioned towards the glass-bottom of Petri dishes (Mattek) to achieve dorsal views of the hindbrain. Imaging was performed at 23°C on a Leica SP8 inverted confocal microscope using a 20× objective (NA 0.7). Software Leica Application Suite X (LAS X) was used to export images in 8-bit format. They were further processed with ImageJ 1.53c.

### Light sheet imaging for cell lineage analysis

Anesthetized life double transgenic Tg[BCP:H2AmCherry;CAAX:GFP] or Tg[BCP:H2AmCherry;HuC:GFP] embryos were mounted individually in 1% LMP-agarose in glass capillaries. During imaging, the embedded embryo was extruded from the capillary into the chamber filled with fish water. Time-lapse imaging was performed at 23°C on a Luxendo (Bruker) MuVi SPIM microscope using 22.2× objective for 16–18 h. Afterwards, the embryonic developmental stage was corrected accordingly (at 23° development is delayed about 0.7-fold). Green and red channels were simultaneously recorded for the entire system every 5 min (Videos 1–2).

### 3D+time cell lineage analysis pipeline

#### Image pre-processing

Image pre-processing was done using Fiji software (NIH). h5 files were converted into 16-bit tiff files using Big Data Processor tool and then cropped to focus on a single hindbrain boundary region. To compensate for small drifts of the embryo during image acquisition, a semi-automated rigid registration was carried out using developed FIJI-scripts (Dyballa et al., 2017). To improve signal to noise ratio on the nuclei BCP:H2AmCherry channel in Tg[BCP:H2AmCherry;CAAX:GFP] embryos, the signal of the plasma membranes (CAAX:GFP) was subtracted and the resulting images were used for single-cell tracking analysis.

#### Cell tracking analyses

For cell-tracking analyses, all boundary cells of the Tg[BCP:H2AmCherry;CAAX:GFP] embryos datasets were reconstructed at single-cell resolution using MaMuT software (Massive Multi-view Tracker software (Wolff et al., 2018). Note that deep- and detailed analyses were performed in one dataset (ID: 1812-r5r6A). Those observations and conclusions were supported with partial analyses of additional datasets of Tg[BCP:H2AmCherry;CAAX:GFP] or Tg[BCP:H2AmCherry;HuC:GFP] embryos time-lapses. Annotated spots were linked and tracked over time from 33.4 hpf (t0) to 44.7 hpf (tf), every 3.5 min, giving a total of 194 timeframes (ID: 1812-r5r6A-1). A total of 120 tracks were followed and classified according to: i) cell proliferative capacity (proliferative when the cell undergoes division, or non-proliferative when the cell does not divide in a period longer than 8 h), and ii) cell fate (progenitor, if the cell localized in the ventricular domain in tf, or neuron, if the cell localized in the mantle zone in tf). The following cell categories were considered: i) non-proliferative progenitors (NP, n = 26), ii) progenitors dividing symmetrically (SyD, n = 24), generating either two progenitors (PP, n = 22/24) or two neurons (NN, n = 2/24); iii) progenitors dividing asymmetrically (AsyD), generating one progenitor and one neuron (PN, n = 30); and iv) committed neurons (CN, n = 20) when a non-dividing cell migrated to the mantle zone during the tracking. Twenty tracks were excluded from the cell fate classification for not allowing a clearly defined localization at tf (ID: 1812-r5r6-2).

Spots within the tracks were then grouped into selections and displayed with different appearances using the 3D-viewer of MaMut software (ID: 1812-r5r6A-2 to 5). Cell lineages and division events were also displayed in a hierarchical way as lineage trees using the Tracking scheme or using the basic graphical functions of R. The numerical features of each single spot (cell), such as spot location, division time, and spot ID, were exported into csv files for further computational analyses.

#### Morphogenetic studies

The plasma membrane channel (CAAX:GFP) was used to model the neural tube volume in a representative fraction of time-frames (45 timepoints, from frame t0 to frame tf, every 17.5 minutes). The dorsal surface of the neural tube facing the ventricle (or ventricular surface) was plotted using Surface tool on Imaris 8.4.2, and the 3D-coordinates were exported in WRL file format to make computations. Data were converted to PLY file format using meshlabserver 2020.07 (Cignoni et al., 2008). Once the triangular mesh read into `R` (R core team 2020) and using packages `rgl` (Adler and Murdoch, 2020) and `Rvcg` (Schlager, 2017), some parts were clipped to obtain the interesting component of the mesh that included the ventricular surface. From this, the vertical sections at coordinate Y=39 were obtained (Figure S3).

#### Cell trajectory analysis

Data on cell positions within the hindbrain were obtained from the csv files generated by MaMut as *X_t_ = (x_t_, y_t_, z_t_) E R^3^*, (*x_t_*: lateral/medial/lateral, *y_t_*: anterior/posterior, and *z_t_*: dorsal/ventral). Coordinates were measured in micrometers. To make it easier to interpret, the averages of each cell’s positions were computed (Figure S3). To compute distances from a given cell at a given time (t) to the ventricular surface, the given ventricular surface was selected for the closest available time; from this, the vertex with minimum Euclidean distance to the cell position was calculated. That minimum distance was taken as the distance from the cell position to the ventricular surface.

### Pharmacological treatments

Tg[BCP:H2AmCherry;HuC:GFP] sibling embryos were treated either with 10 μM of the gamma-secretase inhibitor LY411575 (Stemgent) or DMSO for control. The treatment was applied into the fish water at 28.5°C from either 20 hpf to 26 hpf, or 30 hpf to 36 hpf. After treatment, embryos were imaged in vivo under a Leica SP8 confocal microscope.

### Quantifications and statistical analysis

For assessing the percentage of neurons derived from the boundary cell population (Figure 2C) and the percentage of boundary cells displaying Notch activity (Figure 4C), we quantified the number of cells expressing mCherry with or without GFP in a constant ROI. This ROI covered the whole BCP in every analyzed boundary using Tg[BCP:H2AmCherry;HuC:GFP] or Tg[BCP:H2AmCherry;tp1:d2GFP] embryos (Tables 1, 2). No significant changes were observed when comparing different boundaries from r2/r3 to r5/r6 (Table 2). Thus, for further quantifications we limited our analysis to r4/r5 and r5/r6 and represented r5/r6 in box-plots (Figure 2C, Figure 4C). For quantifying the percentage of neurons derived from the BCP at 48 and 72 hpf (Figure 6D), we immunostained Tg[BCP:H2AmCherry] embryos with anti-Sox2 and anti-HuC and quantified the number of cells expressing mCherry with or without GFP in a constant ROI covering the whole BCP (Table 3). Mean, standard deviation, and standard error of mean are shown in each case using unpaired t-test with Welch’s correction (*p < 0.05, ****p < 0.0001).

## ACKNOWDELGEMENTS AND FUNDING SOURCES

The authors thank Gopi Shah at the Mesoscopic Imaging Facility at EMBL Barcelona for helping with SPIM imaging, Juan Manuel Fuentes at the Scientific-IT (DCEXS-UPF) for helping with computational analysis, and Sebastien Tosi (IRB, Barcelona) and members of the ALMU facility (CRG-UPF, Barcelona) for technical support during image processing. We also thank S Higashijima for kindly providing the Gal4 donor vector plasmid for generating the CRISPR/Cas9 knock-in, and B Link for the tp1:d2GFP construct. We like to thank Laia Subirana and Marta Linares for technical assistance, Gonzalo Ortiz-Alvarez and other current members of the lab for insight and critical discussions, and Isabel Espinosa-Medina and Esteban Hoijman for critical reading of the manuscript. This work was funded by the Agencia Española de Investigación AEI-PGC2018-095663-B-I00 and RED2018-102553-T grants (MICIU-FEDER) to CP. DCEXS-UPF is a Unidad de Excelencia María de Maeztu funded by the AEI (CEX2018-000792-M). CEP is a recipient of a predoctoral FPU fellowship from the Spanish Ministry of Universities. FU work was supported by PGC2018-101643-B-I00 (MICIU-FEDER). CP is a recipient of ICREA Academia award (Generalitat de Catalunya).

## AUTHOR CONTRIBUTIONS

CFH and CP contributed to the concept and design of experiments, interpretation of the results and wrote the manuscript. CFH and CEP performed and analyzed the experiments and interpreted the results. FU collaborated with CFH in the mathematical analysis of data.

## VIDEOS

**Video 1: Time-lapse of the boundary cell population undergoing neurogenesis.** Tg[BCP:H2AmCherry;HuC:GFP] embryo displayed as a transverse view, with mCherry in the boundary cell nuclei and GFP in the differentiated neurons. Imaging was performed from 32 to 49 hpf. The movie shows the dynamics and behavior of the cells of a single boundary (r5/r6), and the insert is a zoom on the green framed region. A single progenitor cell (white circle) undergoing asymmetric division is tracked upon time. White arrowhead indicates the progenitor cell immediately before division (34.5 hpf), and blue and green arrowheads the progenitor and neuronal derivatives, respectively. Both, progenitor (P) and neuronal (N) fated daughter cells were tracked after division (white circle). Blue and green lines represent the distinct cells trajectories. Note how the neuronal derivative migrates towards the HuC domain and eventually differentiates into a neuron. Right-hand site inserts are dorsal views of that very same boundary cells with anterior to the top, focusing at different Z-positions, corresponding either to the dorsal progenitor derivative (top) or the ventral neuronal derivative (bottom), with tracked cells encircled in white. A, anterior; D, dorsal; M, medial; hpf, hours-post-fertilization. Embryo dataset ID: 0310-r5r6A-1.

**Video 2: Time-lapse of the boundary cell population.** Tg[BCP:H2AmCherry] embryo displaying mCherry in the boundary cell nuclei was imaged from 32 to 44.7 hpf. The movie shows the dynamics and behavior of a single embryo with all boundaries, and the insert focusses on a single boundary. Dorsal view with anterior to the top, with inserts of r4/r5 as indicated by the green box. Green arrowheads in the insert indicate cell division events. Transverse view of r5/r6 boundary. A, anterior; D, dorsal; M, medial; BCP, boundary cell population; hpf, hours-post-fertilization. Embryo dataset ID: 1219.

## SUPPLEMENTARY MATERIAL

**Supplementary Figure S1: Generating CRISPR/Cas9 knock-in transgenic lines to trace the boundary cell population (A) Scheme depicting the CRISPR/Cas9 knock-in strategy.** The boundary cell population cis-regulatory element (BCP-CRE) located upstream sgca transcription start site was targeted by CRISPR/Cas9; after inducing double-strand breaks, the donor vector carrying Gal4FF was integrated. Using the hsp70 promoter in the construct allowed both forward and reverse integrations to be functional. (B) Genomic description of the knock-in target site, before and after integration and repair, in one of the Tg[BCP:Gal4] founders (the one selected for this work). The sgRNA target site is indicated in blue; intergenic sequence, grey; and sgca, black (the coding region of the first exon is underlined). After efficiently knocking-in the target locus, two copies of the insert were integrated in tandem (orientation indicated with purple boxes; green sequence indicates a fraction of Gbait, and purple sequence the beginning of hsp70 promoter). Note the presence of small indels (grey sequence between the blue sequence of the endogenous locus and the integrated construct) indicating that homology-independent DNA repair occurred in the targeted locus. (C) CRISPR/Cas9 knock-in pipeline. Three sgRNA target sites (#1, #2, #3) were designed according to the best hits provided by the crisprscan.org and chopchop.cbu.uib.no platforms; these were synthetized, and their efficiency was confirmed by detecting indels in the genome target site. Tg[UAS:KAEDE] embryos at the one-cell stage were injected with a mix containing the two sgRNAs (sgRNA #2 for digesting genomic DNA, and Gbait sgRNA for digesting the donor plasmid), Cas9 protein and the donor DNA vector. Embryos were selected by mosaic KAEDE-expression (53% efficiency, n = 40/76). Successful Gal4 genome integration was confirmed in adult fish by PCR with Gal4FF primers and resulted in 59% efficiency (n = 16/27). Knock-in founders were selected by KAEDE-expression in F1 offspring and displayed a germline transmission rate of 25% (n = 4/16). 75% of these founders specifically expressed KAEDE in the boundaries (n = 3/4). Finally, adults were genotyped by fin clip using a strategy that confirmed integration and the insert directionality; this involved a combination of PCRs using Fw1, Rv1, Fw2 and Rv2 primers, so that the products A, B, C or D were amplified depending on the construct orientation. The selected Tg[BCP:Gal4] fish used in this work showed a A+D+ genotype, indicating that it contained at least two copies of the donor vector inserted in tandem (forward and reverse). Subsequent sequencing of genomic DNA (gDNA) target site confirmed the tandem knock-in (see the sequence description in (B)). The Tg[BCP:Gal4;UAS:KAEDE] line (called Tg[BCP:KAEDE] in the manuscript) was outcrossed to segregate the UAS:transgene and to isolate Tg[BCP:Gal4]. After this, several transgenic lines were generated by crossing Tg[BCP:Gal4] fish with different UAS:reporter (UAS:GFP, UAS:H2AmCherry).

**Supplementary Figure S2: Comparative analyses of the spatiotemporal profile of the boundary cell population markers and the Tg[BCP:H2AmCherry] transgene** (A–F) Double in situ hybridization of Tg[BCP:H2AmCherry] embryos at different stages with sgca and mCherry. Dorsal, transverse, or sagittal views are indicated for each case. Expression of mCherry was delayed with respect to sgca expression. (G–N) In situ hybridization with the boundary marker rfng in Tg[elA:GFP] embryos (a landmark of r3 and r5) at different stages, combined with the immunostaining of ZO-1, to label apical tight junctions. Note the offset of rfng between 31 and 33 hpf (M, N). (O, P) Double immunostaining of wild-type embryos at 30 hpf with Taz and ZO-1. Taz is expressed in the ventricular domain of the boundaries (Voltes et al., 2019); ZO-1 labelling allows the larger apical feet of boundary cells to be visualized (Symonds et al., 2020). Q-R) Transverse views of the Tg[BCP:H2AmCherry;4xGTIIC:d2GFP] embryos showing that mCherry-boundary cells display Yap/Taz-TEAD-activity at 33 hpf (Q-Q’) and 48 hpf (R-R’). Inserts show Yap/Taz-TEAD active boundary cells (black asterisks) in the progenitor domain at 33 hpf and a fraction of their derivatives in the neuronal differentiated domain at 48 hpf (white asterisks). White arrowheads indicate the position of the hindbrain boundaries. BCP, boundary cell population; hpf, hours-post-fertilization; MHB, mid-hindbrain boundary. Scale bar, 50 μ

**Supplementary Figure S3: Temporal representation of morphogenetic and cellular movements in the boundaries** (A) Dynamic comparison of the segmented ventricular surface of rhombomere 5 over time, displaying it colored at the indicated time on the top of the 45 different analyzed time frames (in grey hues). (B) Overlay of segmented ventricular surfaces of the same embryo over time, color-coded according to time of development. Note the flattening that it undergoes as the neural tube grows, and the shift towards dorsal. (C) Average movement of all boundary progenitor cells along the dorsoventral axis over time, measured by the μm of distance to the ventricular surface. Each frame corresponds to the average position of all individual progenitors. Note that the progenitors undergo a shift on their position towards dorsal (e.g., closer to the ventricular surface), coinciding in time with the increase in the neuronal differentiated population located in the ventral domain. (D) Cell movements along the lateromedial axis over time. The position of each single progenitor cells was tracked over time (black lines). Red lines represent the average position for the tracks grouped by hemisphere. Note that no relevant cellular movements were observed along this axis. D, dorsal; V, ventral; L, lateral; M, medial; hpf, hours-post-fertilization.

**Supplementary Figure S4: Expression of Notch-players in the hindbrain** (A, B) Tg[BCP:H2AmCherry;tp1:d2GFP] embryos in vivo imaged (A) or immunostained with the radial glia marker zrf1 (B) at 48 hpf. Transverse views with inserts showing the cell processes of Notch-active boundary cells towards the basal domain of the hindbrain (A), and equivalent Notch-active cells stained with anti-zrf1 (B, blue). Note that Notch-active boundary cells display radial glia morphology. (C–H) Embryos displaying Notch-activity and Notch players in the rhombomeres at 26 and 36 hpf. (C, E) Tg[tp1:d2GFP] as a read-out of Notch activity. (D–D’, F–F’, G) Embryos hybridized with notch3 and deltaD. (H–H’) Embryos hybridized with notch1. Transverse views are shown at the level of rhombomere 5. BCP, boundary cell population; hpf, hours-post-fertilization. Scale bar, 50 μ

