## Supplementary figures and images for "The neurogenic fate of the hindbrain boundaries relies on Notch-dependent asymmetric cell divisions"

### Figure S1

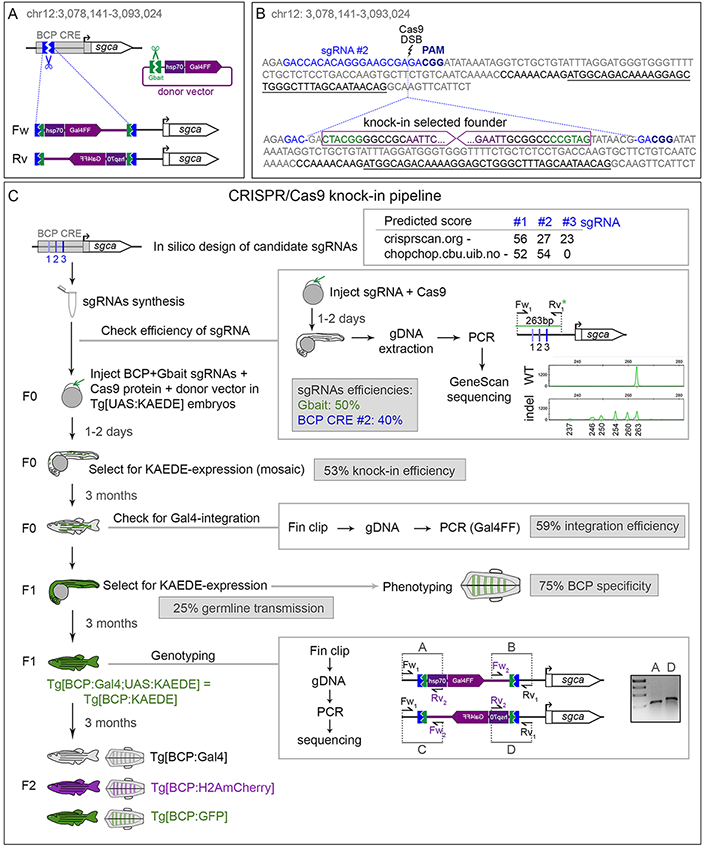

### Figure S2

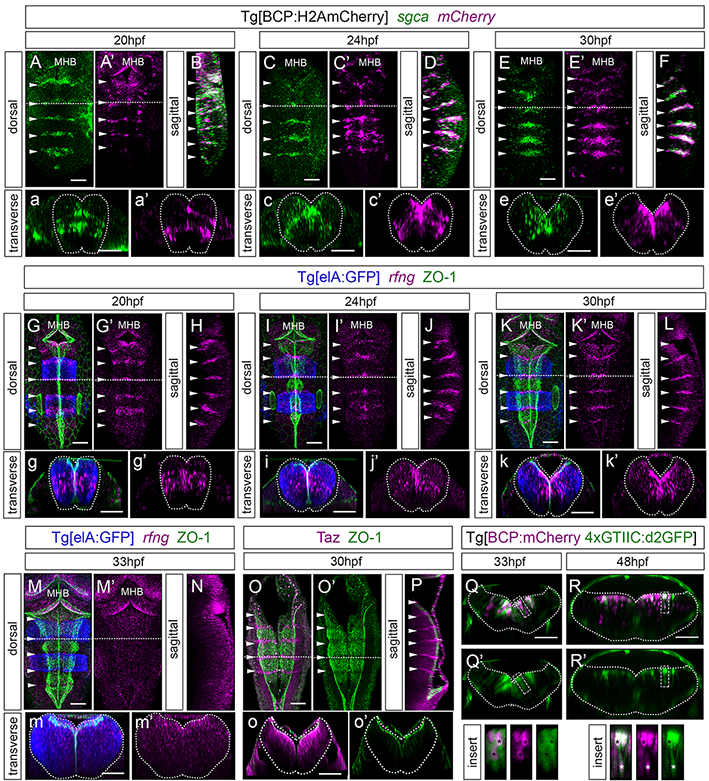

### Figure S3

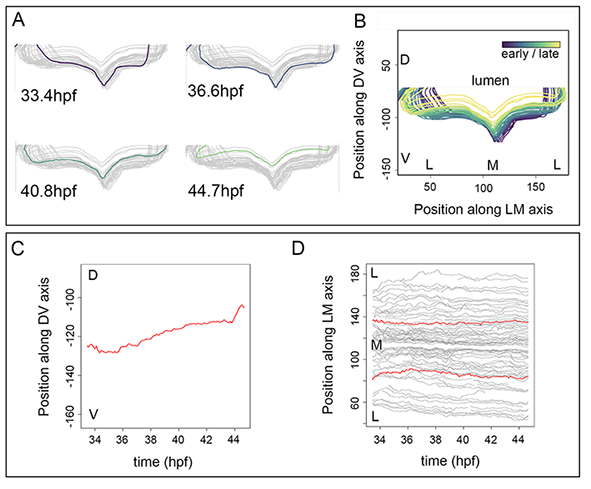

### Figure S4

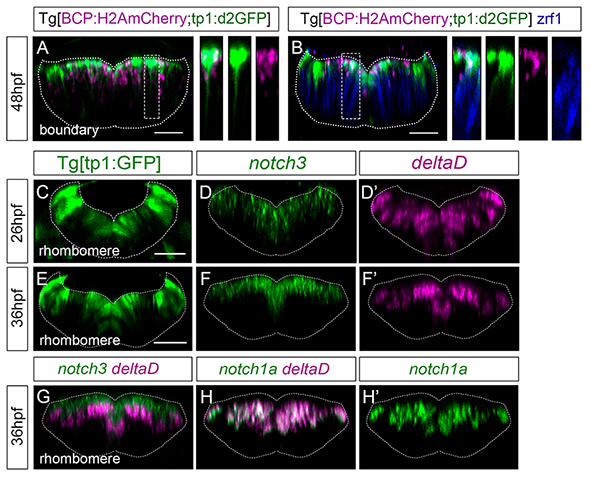
